# Ewing Sarcoma Breakpoint Region 1 (EWSR1) protein binds to the non-structural protein-2 and the non-coding regions of the Chikungunya virus genome to inhibit the virus replication

**DOI:** 10.64898/2025.12.16.694598

**Authors:** Shivani Balyan, Anshula Sharma, Devendra Sharma, Deepti Jain, Sudhanshu Vrati

**Affiliations:** Regional Centre for Biotechnology, Faridabad-121001, India

**Keywords:** Alphavirus, EMSA, BLI, NCR, RNA-protein binding

## Abstract

Chikungunya virus (CHIKV) is an alphavirus with a single-stranded RNA genome. Conserved sequence elements within the 5′- and 3′-non-coding regions (NCRs) of the CHIKV genome interact with host proteins to regulate viral RNA replication. Electrophoretic mobility shift assays (EMSA) and mass spectrometry identified several host RNA-binding proteins that associate with these conserved NCR elements, among which EWSR1 uniquely bound the 3′-ends of both the plus-sense genomic RNA and the minus-sense replication intermediate RNA. EMSA and biolayer interferometry demonstrated that EWSR1 binds strongly to two RNA elements: a 48-nucleotide region in the plus-strand 3′-NCR (3NCR48) and a 50-nucleotide region at the 3′-end of the minus-strand replication intermediate (5NCR-RC50), which is complementary to the genome’s 5′-NCR and its adjacent sequence. RNA immunoprecipitation from CHIKV-infected cells confirmed that EWSR1 binds CHIKV RNA *in vivo*. Functional analyses revealed that CHIKV infection downregulates EWSR1 expression and causes its cytoplasmic accumulation. Knockdown or knockout of EWSR1 expression increased CHIKV genomic RNA levels and viral titers, whereas its overexpression suppressed both. EWSR1 reduced the synthesis of both plus- and minus-sense CHIKV RNAs in infected cells. Additionally, EWSR1 was found to bind CHIKV nsP2 and inhibit its helicase activity, which is critical for viral genome replication. The antiviral activity of EWSR1 extended to Ross River virus but not to Sindbis virus, consistent with conservation of the 48-nucleotide sequence in their respective 3′-NCRs. Together, these findings identify EWSR1 as a novel host restriction factor that limits CHIKV replication through specific interactions with viral RNA and the nsP2 protein.

**IMPORTANCE:** Chikungunya virus (CHIKV) poses a significant medical challenge, necessitating the development of novel antiviral strategies. This study demonstrates that the host protein EWSR1 plays a key role in controlling CHIKV replication. By binding to specific regions of viral RNA and interacting with nsP2, a critical viral enzyme, EWSR1 inhibits viral RNA synthesis and reduces viral titers. In the absence of EWSR1, CHIKV replicates more efficiently, underscoring its antiviral function. These findings establish EWSR1 as a novel host restriction factor that limits CHIKV replication. Understanding this interaction enhances our knowledge of host–virus dynamics and may guide the design of new antiviral approaches against CHIKV and related alphaviruses.

## INTRODUCTION

Chikungunya virus is a single-stranded plus-sense RNA genome containing enveloped virus. It belongs to the Alphavirus genus of the *Togaviridae* family and is transmitted by the *Aedes* mosquitoes, primarily *Aedes aegypti* and *Aedes albopictus*. The virus causes Chikungunya fever, characterized by maculopapular rash, myalgia, arthralgia, nausea, and chills (1). CHIKV has caused periodic epidemics in Africa and Asia from the 1960s to the 2000s, a large-scale epidemic in the Indian Ocean region in 2005–2006, and a major outbreak in India in 2006 with 1.25 million cases. The virus has continued to cause frequent outbreaks across Asia, Africa, and the Americas, spreading to over 110 countries worldwide. In 2024 and as of 30 November, approximately 480,000 CHIKV cases and over 200 deaths were reported globally (2). Moreover, the virus causes severe, long-lasting indisposition in a large population each year. While a couple of CHIKV vaccines have recently been licensed (3–5), no antivirals are currently available. Understanding the virus replication and identification of critical events, such as the viral RNA interaction with the host protein/s, will greatly help in designing novel antivirals.

CHIKV has a ∼11.8 kb genome having a 5’-methylguanylated cap and a 3’-polyA tail. The genome encodes four non-structural and five structural proteins. The non-structural proteins (nsP1, nsP2, nsP3, and nsP4) are synthesised from two-thirds of the genome towards its 5’-end. They are components of the replication complex required for genomic RNA (1) replication. The structural proteins (C, E3, E2, 6K, and E1) are synthesised from the sub-genomic RNA, comprising one-third of the genome towards the 3’-end (1). The CHIKV virion consists of an icosahedral nucleocapsid composed of the capsid protein and genomic RNA surrounded by the lipid envelope containing the E1 and E2 glycoproteins (6). While E1 induces pH-dependent fusion with the endosomal membrane, E2 protein binds to cell surface attachment receptors (7).

The open reading frames in the CHIKV genome are flanked by 60- and 529-nucleotide non-coding regions (NCRs) at the 5′- and 3′-ends of the genome, respectively. The alphavirus NCRs contain conserved sequence elements (CSEs) that play an important role in the viral life cycle and the synthesis of the minus- and plus-sense viral RNAs (8). These NCRs interact with the viral and host cell proteins and facilitate viral replication. A highly conserved AU dinucleotide at the very beginning of the 5′-NCR is essential for genome replication, likely serving as a recognition site for the viral RNA-dependent RNA polymerase (RdRp) during initiation of the plus-sense RNA synthesis (9–11). The 5′-NCR contains a stem-loop structure with a role in the minus-strand RNA synthesis replication (9–11). Besides, it contains core promoter elements necessary for the initiation of both minus- and plus-sense RNA synthesis along with other cis acting RNA elements and 3’UTR (11, 12). These elements are critical for the recruitment of the viral RdRp and other replication factors, including the viral and host proteins. Near the start of the nsP1 coding region (just downstream of the 5′-NCR), a 51-nucleotide CSE is present, which is important for the virus replication, especially in mosquito cells (11).

The 3′-NCR also contains CSEs necessary for efficient viral RNA replication (11–13). Immediately upstream of the poly(A) tail, a short, highly conserved sequence (19–24 nucleotides) is present in all alphaviruses. This element is essential for viral replication and acts as a core promoter for the minus-sense RNA synthesis (14). The 3′-NCR contains multiple short repeat sequence elements (RSEs), which are often arranged in tandem and vary in copy number among different alphavirus species. These RSEs bind host proteins and are important for efficient virus replication (11, 12, 14). A U-rich sequence of ∼60 nucleotides is often found upstream of the conserved 19-nucleotide region. This domain interacts with host proteins to stabilise the viral genome (14). Recent studies have identified a highly conserved Y-shaped stem-loop structure within the CHIKV 3′-NCR that plays roles in replication efficiency and possibly in evading host defences (15).

Several host proteins have been identified that bind the alphavirus NCRs. The 5′-NCR interacts with host translation initiation factors eIF4E and eIF4F, which are critical for initiating protein synthesis (11). Heterogeneous nuclear ribonucleoprotein A1 (hnRNP A1) binds with the 5′-NCR of Sindbis virus (SINV) RNA and enhances the synthesis of minus-sense RNA (16). The cellular HuR protein that binds with the U-rich elements of the 3’-NCR of SINV RNA, and upstream of the 3’-CSE in the 3’-NCRs of Ross River virus (RRV) is important for the viral genome stability and its efficient replication (11, 17, 18). The La autoantigen also interacts with the 3′-NCR, likely contributing to RNA stability and possibly influencing translation or replication (11, 19).

Collectively, these studies illustrate the pivotal role of CSEs and their interaction with host proteins in alphavirus replication. In the present work, several host proteins that specifically recognize CSEs in the CHIKV NCRs have been identified, with a focus on determining the functional impact of EWSR1 in CHIKV replication.

## MATERIALS AND METHODS

### Cells and viruses

Human embryonic kidney cells (HEK293T) and Embryonal rhabdomyosarcoma (ERMS) cells were grown in Dulbecco’s modified Eagle’s medium (DMEM) (Himedia, AL007A) with 10% foetal bovine serum (FBS) (Gibco, 10270106). The African green monkey kidney cells (Vero) and baby hamster kidney cells (BHK-21) were cultured in Minimum essential medium (MEM) (Himedia, AL0475) with 10% FBS. HAP1 cells (a near-haploid human cell line derived from the KBM-7 chronic myelogenous leukemia cell line) were grown in Iscove’s modified Dulbecco’s medium (IMDM) (HiMedia, AL070S) with 10% FBS. All cells were grown at 37 °C in a humidified atmosphere with 5% CO_2_. All cultures were supplemented with 1× Penicillin-Streptomycin-Glutamine solution (HiMedia, A001A). The IND-06 Guj strain of Chikungunya virus (CHIKV) (GenBank accession number JF270482.1), EG AR 1019 strain of Sindbis virus (SINV) and T48 strain of Ross River virus (RRV) (GenBank accession number GQ433359.1) were used in this study. The IND-06 Guj strain of CHIKV was isolated in the lab from a CHIKV patient of the 2006 Gujarat outbreak. SINV and RRV were obtained from the World Reference Centre for Emerging Viruses and Arboviruses, University of Texas Medical Branch, USA. The viruses were grown in BHK-21 cells and their titers determined by plaque assay.

### Plaque assay

Vero cells were seeded in 6-well tissue culture plates overnight to achieve monolayers with approximately 80% confluency. The cells were infected with the serial dilutions of virus-containing culture supernatants prepared in the incomplete MEM. After 1 h incubation, the cells were washed with 1× PBS and incubated at 37 °C with an overlay of 2% low-melting agarose (Sigma Aldrich, A4018) and equal volume of 2× MEM supplemented with 10% FBS and 1× Penicillin-Streptomycin-Glutamine solution (HiMedia, A001A) for 36 h. The cell monolayers were then fixed with 4% formaldehyde and stained with 0.2% crystal violet to visualize the plaques.

### Electrophoretic mobility shift assay (EMSA)

EMSA was performed as described previously with slight modifications (20). The HEK293T, Vero, and ERMS cells were harvested and lysed using the lysis buffer containing 25 mM Tris pH 7.4, 150 mM NaCl, 1 mM EDTA, 1% NP-40, and 5% glycerol. The protease inhibitor cocktail (Sigma Aldrich, P8340) was added immediately before cell lysis. The lysates were incubated on ice for 1 h with periodic vortexing and spun down, and the supernatant was collected for protein estimation by the Bradford Assay kit (Thermo Scientific, 23200). Ten μg cell lysate was incubated with 10 μg poly(I)-poly(C) (polyIC) (Merck, P1530), 20 ng biotin-labelled synthetic RNA, and 1 U RNasin (Promega, N2611) in the binding buffer (14 mM HEPES pH 7.5, 6 mM Tris-Cl pH 7.5, 1 mM EDTA, 1 mM DTT, and 60 mM KCl) in a final volume of 25 μl for 30 min at room temperature with shaking. The RNA-protein mixture was then crosslinked using a UV torch (4 watts) for 30 min. The RNA-protein complexes were electrophoresed on a nondenaturing 12% polyacrylamide gel (acrylamide/bisacrylamide ratio 29:1) at 4 °C in 0.5X Tris-borate-EDTA buffer. The gel was transferred to a Biodyne nylon membrane (Thermo, 77016) at 200 mA, 4 °C for 2 h. The membrane was then developed using a chemiluminescent nucleic acid detection module kit as per the manufacturer’s protocol (Thermo, 89880), in which the blot was blocked and subjected to streptavidin-HRP conjugate, and then HRP was detected using luminol enhancer and visualised on X-ray film.

### RNA pull-down and mass spectrometry

Two mg cell lysate was incubated in binding buffer (14 mM HEPES pH 7.5, 6 mM Tris-Cl pH 7.5, 1 mM EDTA, 1 mM DTT, and 60 mM KCl) with 2 mg poly(I)-poly(C), 4 μg synthetic RNA and 1 U RNasin in a final volume of 2 ml for 45 min at room temperature with shaking. After this, the RNA-protein complexes were crosslinked with a UV torch for 30 min. Dynabeads MyOne streptavidin C1 (Invitrogen, 65001) beads were washed thrice with binding buffer. RNA-protein mix was added to the beads and incubated for 30 min at room temperature with shaking. Beads were then washed with binding buffer, and bound proteins were eluted using 95% formamide and 10 mM EDTA at 65 °C for 5 min. The eluted proteins were then buffer exchanged with 50 mM ammonium bicarbonate, digested with 1 μg trypsin (Promega, V5280) at 37 °C for 18 h, and analyzed by mass spectrometry (SCIEX Triple TOF 5600+).

### siRNA transfection and virus infection of the cells

ON-TARGETplus SMARTpool small interfering RNAs (siRNAs) targeting EWSR1 and nontargeting control were obtained from Dharmacon. HEK293T cells were transfected with 30 nM siRNA using DharmaFECT1 (Horizon Discovery, T-2005-01) following the manufacturer’s instructions. At 48 h post-transfection, cells were infected with CHIKV, RRV, or SINV and harvested at different times post-infection (pi). Cells and supernatant were subsequently processed for western blotting, qRT-PCR, and plaque assays, respectively.

### Generation of the knockout (KO) HAP1 cells

HAP1 cells were transfected with the CRISPR/Cas9 EWSR1 knockout plasmid synthesized by GeneScript. The knockout plasmid was based on GenScript plasmid eSpCas9-2A-Puro (PX459) V2.0. The sgRNA sequence (sgEWSR1: 3’-TACACCGCCCAGCCCACTCA-5’) was selected from the GeCKO v2 Human CRISPR Knockout Pooled Library and reviewed by the CHOPCHOP web tool. The sgRNA/Cas9 plasmid was transfected using Lipofectamine 3000 (Thermo, L3000001). The transfected cells were selected using puromycin solution (InvivoGen; ant-pr-1) at a 0.8 µg/mL concentration. The absence of the EWSR1 expression in the KO cells was confirmed by western blotting.

### EWSR1 ectopic expression studies

The full-length EWSR1 cDNA was amplified using total cellular RNA and the following synthetic oligonucleotide primers: forward primer-ATGC<u>GTCGAC</u>ATGGCGT CCACGGAT (Sal I restriction site underlined) and reverse primer-ATAT<u>GCGGCCGC</u>CTAG TAGGGCCGATCT (Not I restriction site underlined). The cDNA was cloned into the pCMV Script vector (Novopro, V011243) under the CMV promoter. HEK293T cells were transfected with the plasmid using Lipofectamine 3000 according to the manufacturer’s protocol. At 24 h post-transfection, cells were infected with CHIKV, RRV or SINV and harvested at 12 h post-infection. Cells and supernatant were then processed for western blotting and qRT-PCR, and plaque assay, respectively.

### Nuclear and cytoplasmic fractionation of cell lysate

HEK293T cells were infected with CHIKV and harvested at 12 h pi. The nuclear and cytoplasmic fractions were obtained using the NE-PER nuclear and cytoplasmic extraction reagent (Thermo Fisher, 78833) as per the manufacturer’s protocol. GAPDH and Lamin A/C were used to establish the purity of cytoplasmic and nuclear fractions, respectively.

### RNA isolation and quantitative real-time PCR

The total RNA was isolated from the cells using the RNAiso Plus reagent (Takara, 9109) and purified using the phenol-chloroform method. The first strand of the cDNA was synthesized using 100 ng total RNA with the ImProm II Reverse Transcriptase kit (Promega, A3802). The quantitative real-time PCR (qRT-PCR) was set up using the TB Green Premix Ex Taq (Takara; RR420A). The reactions were performed using QuantStudio 6 (Applied Biosystems). The following primers were used: CHIKV forward-AAGCTYCGCGT CCTTTACCAAG, CHIKV reverse CCAAATTGTCCYGGTCTTCCTG; SINV forward-AAAGGATACTTTCTCCTCGC, SINV reverse-TGGGCAACAGGGACCATGCA; RRV forward-GCGACGGTGGATGTCAAGGAG, RRV reverse-AGCCAGCCCACCTAACCCACTG; GAPDH forward-GGTGAAGGTCGGAGTCAACG, GAPDH reverse-AGGGATCTCGCTCCTGGAAG; EWSR1 forward-GGGTATGGCACTGGTGCTTAT, EWSR1 reverse-CAGACTGAGCTGCATAGGAGG.

### RNA immunoprecipitation

HEK293T cells were infected with CHIKV and harvested at 12 h pi. RNA immunoprecipitation (RIP) was performed using a MAGNA RNA Binding protein RIP kit (Sigma Aldrich, 17-700) according to the manufacturer’s instructions. Cells were lysed in RIP lysis buffer, and an equal amount of lysate was incubated with either EWSR1 antibody or an isotype-matched IgG antibody bound to protein A/G magnetic beads. After washing, RNA-protein complexes were treated with proteinase K to digest the protein, and RNA was purified using the phenol-chloroform method. qRT-PCR to determine the CHIKV RNA levels was performed as described earlier.

### Recombinant EWSR1 protein expression and purification

Full-length EWSR1 cDNA was amplified using total cellular RNA and the following synthetic oligonucleotide primers: forward primer-ATGCCG<u>GAATTC</u>GCGTCCACG GATTAC with EcoRI site (underlined), reverse primer-ATAT<u>GCGGCCGC</u>CTAGTAGGG CCGATCTC with NotI site (underlined). The cDNA was cloned under the T7 promoter into the pET28a (Novagen, 69864) vector. *Escherichia coli* BL21-DE3 cells transformed with the pET28a-6xHis-EWSR1 plasmid were grown in Luria Broth at 37 °C until absorbance A_600_ reached 0.6-0.8. The recombinant protein expression was then induced by the addition of 0.5 mM isopropyl-β-d –thiogalactoside (IPTG), followed by incubating the cells under shaking at 22 °C for 14 h. For the purification of 6xHis-tagged EWSR1 protein, cell pellets were centrifuged at 4,000 g at 4 °C for 20 min and resuspended in lysis buffer containing 50 mM Tris–HCl pH8.0, 500 mM NaCl, 5 mM EDTA, 1 mM PMSF and then lysed by sonication (Sonics and materials, 13mm probe, 55% amplitude, 10 cycles with pulses of 10 sec on and 50 sec off). The cell lysates were centrifuged at 16,000 g for 20 min at 4 °C. The pellet was resuspended in wash buffer A (50 mM Tris-Cl pH 8, 500 mM NaCl, 1% Triton X-100, 5 mM EDTA, 1 mM PMSF) and then sonicated again. The cell lysates were centrifuged at 16,000g for 20 min at 4 °C and the pellet was homogenised in wash buffer B (50mM Tris-Cl pH8, 500mM NaCl, 1mM PMSF). This was followed by centrifugation at 16,000 g for 20 min at 4°C. The pellet was then solubilised in 50 mM Tris-Cl pH 12, 500 mM NaCl, 5 mM EDTA, 1 mM PMSF, 2 M urea and incubated at room temperature for 1 h with periodic vortexing, and then centrifuged at 16000 g for 30 min. Supernatant containing EWSR1 was then allowed to refold in 50 mM Tris-Cl pH 8, 10% sucrose, 1 mM EDTA, 1 mM PMSF for 14 h at 4 °C. The refolded sample was then centrifuged at 16000 g for 30 min. Supernatant was then loaded onto a pre-equilibrated Ni-NTA column, washed with washing buffer containing 20 mM Tris–HCl pH 8.0, 500 mM NaCl, and 20 mM imidazole, followed by elution with the elution buffer containing 20 mM Tris-HCl pH 8.0, 500 mM NaCl, and 200 mM imidazole (21). The eluted protein was quantified by the Bradford Assay kit (Thermo Scientific, 23200). The protein identity and purity were established by electrophoresis on a 12% SDS-PAGE followed by western blotting, as well as by peptide mass fingerprinting.

### Expression and purification of CHIKV nsP2 protein

The full-length CHIKV nsP2 protein, and its helicase and protease domains were expressed and purified as described previously (22). Briefly, a modified pET14b expression plasmid encoding the gene of interest and an N-terminal His_6_-SUMO tag was used to transform *Escherichia coli* Rosetta (DE3) cells. Cultures were grown in LB to an OD_600_ of 0.6–0.8 and induced with 0.5 mM Isopropyl β-d-1-thiogalactopyranoside (IPTG) at 18 °C for 16 h. Cells were harvested and lysed in buffer containing 20 mM HEPES pH 7.5, 500 mM NaCl, 5% Glycerol and 2 mM β-Mercaptoethanol. The lysate was clarified and applied to a Ni-NTA column (Cytiva), followed by elution using an Imidazole gradient. His_6_-SUMO tag was removed using PreScission protease, and the sample was dialysed overnight at 4°C. The protein was concentrated and subjected to size-exclusion chromatography using the Superdex 200 16/600 (Cytiva) column in 20 mM HEPES pH 7.5, 300 mM NaCl, 5% Glycerol, and 0.5 mM DTT. The purity was confirmed by SDS PAGE.

### Western blotting

The cells were harvested and lysed using the lysis buffer containing 25 mM Tris pH 7.4, 150 mM NaCl, 1 mM EDTA, 1% NP-40, and 5% glycerol. The protease inhibitor cocktail (Sigma Aldrich, P8340) was added immediately before cell lysis. The lysates were incubated on ice for 1 h with periodic vortexing and spun down, and the supernatant was collected for protein estimation by the Bradford Assay kit (Thermo Scientific, 23200). The 5× SDS-PAGE loading dye was then added to the lysates and heated at 95 °C for 10 min. Equal amounts of protein lysates were then resolved on a 12% SDS-PAGE and transferred onto a polyvinylidene difluoride (PVDF) membrane. The membrane was then blocked with 5% BSA for 1 h at room temperature, followed by overnight incubation with the primary antibody at 4 °C. The membrane was then washed with PBS with 0.1% Tween-20 and incubated with horseradish peroxidase (HRP)-conjugated secondary antibody for 1 h. The protein bands were visualized using the SuperSignal West Pico PLUS Chemiluminescent substrate (Thermo Scientific; 34577).

### Biolayer interferometry (BLI)

EWSR1 binding to RNAs was studied by the ForteBio Octet Red instrument. The ligand, biotinylated-RNA (50 ng) in nuclease free water was loaded on the surface of a streptavidin-conjugated biosensor for 600 sec. The binding assays were performed in black 96-well plates at 25 °C, under agitation (1000 rpm), in 200 µl binding buffer (14 mM HEPES pH 7.5, 6 mM Tris-Cl pH 7.5, 1 mM EDTA, and 60 mM KCl). Initially, a 120 sec biosensor baseline step was performed in buffer only. Following this, association of the biosensor-bound RNA ligand and the EWSR1 protein (0-800 nM) in the binding buffer was recorded for 180 sec. Finally, dissociation was followed for 180 sec. For the CHIKV nsP2 and EWSR1 interaction, histidine-tagged EWSR1 (10 µg) was loaded on the surface of a Ni-NTA biosensor for 300 sec. The sensor loaded with EWSR1 was allowed to associate with CHIKV nsP2 (100-1000 nM) for 400 sec in 50 mM Tris buffer (pH 7.4). The dissociation was then followed for 800 sec. Correction of any systematic baseline drift was done by subtracting the shift recorded for the sensor loaded with the ligand in the absence of analyte. The BLI sensorgram was plotted showing the biosensor’s response against time for different analyte concentrations. The data were fitted to a 2:1 binding model to determine the equilibrium dissociation constant (*K*_d_). Only data with an R^2 value of >0.95 were included in the analysis.

### Strand-specific quantitation of CHIKV RNA

The plus- and minus-sense RNA of CHIKV were quantified using a strand-specific qRT-PCR assay as described previously (22). Briefly, the total RNA isolated from the CHIKV-infected cells was employed for the cDNA synthesis by reverse transcription. Tagged (non-viral sequence) primers PtagCHIKp and NtagCHIKn were used for the cDNA synthesis from the plus- and minus-sense CHIKV RNA, respectively. The real-time qPCR was performed using a combination of primers that bind to the non-viral tag sequence and viral strand. The strand-specific RNA copy numbers were determined using the plus- and minus-sense RNA-specific standard curves.

### Co-immunoprecipitation

HEK293T cells were infected with CHIKV for 12 h. The cells were washed with 1× PBS and lysed using RIPA lysis buffer. The CHIKV proteins were co-immunoprecipitated from 1 mg cell lysate and 5 µg anti-EWSR1 antibody using the Co-immunoprecipitation kit (Thermo Scientific; 26149). The immunoprecipitated proteins were western blotted using the CHIKV anti-nsP2 antibody (Genetex; 636897).

### Helicase assay

To investigate the effect of EWSR1 on RNA unwinding activity of CHIKV nsP2, a helicase assay was performed following the protocol described previously (23, 24). The assay utilised an Alexa Fluor 488-labelled 28-mer ssRNA oligonucleotide (Alexa488-ssRNA, 5’-AAAAAAAAAAAACCAGGCGACAUCAGCG-3’) and an unlabelled 16-mer ssRNA oligonucleotide (5’-CGCUGAUGUCGCCUGG-3’) to generate a dsRNA helicase substrate with a 12-base 5’ overhang. To prepare the dsRNA substrate, the labelled and unlabelled RNA oligonucleotides were mixed at a 1:1.1 molar ratio in a buffer containing 10 mM HEPES (pH 7.2) and 20 mM KCl. To facilitate annealing, the RNA mixture was heated to 95 °C for 1 min in a thermal cycler, followed by gradual cooling at a rate of 1 °C per min until reaching 22 °C. The optimized reaction mixture for the unwinding assay consisted of the strand-displacement assay buffer (40 mM HEPES, pH 7.5, 2 mM dithiothreitol (DTT), and 12 mM NaCl), 1 μM purified CHIKV nsP2 protein, 50 nM dsRNA substrate, 800 nM unlabelled RNA trap (5’-CCAGGCGACAUCAGCG-3’), 20U RNaseOUT inhibitor (Thermo Fisher Scientific, 10777019), and varying concentrations of EWSR1. The reaction mixture was incubated at 26°C for 20 min in a thermal cycler to facilitate the formation of the protein RNA complex. Subsequently, a 3.5 mM ATP-magnesium acetate mixture was added, followed by incubation at 37 °C for 120 min. The reactions were then terminated by adding a stop solution containing 100 mM Tris-HCl, pH 7.5, 0.1% bromophenol blue, 1% SDS, 50 mM EDTA, and 50% Glycerol. The separation of ssRNA and dsRNA was carried out through electrophoresis on a 15% nondenaturing polyacrylamide gel at 4°C. The RNA bands were visualized using a Typhoon imager (Cytiva), and their intensities were quantified using the ImageJ software.

### CHIKV nsP2 protease assay

The proteolytic activity of CHIKV nsP2 protease was determined using a FRET-based approach (22, 25, 26). The assay was performed *in vitro* using the purified CHIKV nsP2 protein and the FRET-based octapeptide substrate {DABCYL}-RAGG↓YIFSS-{Glu(EDANS)}-NH2 (Biolink) representing the nsP3/4 site (27). For the assay, 1 μM CHIKV nsP2 was incubated with different concentrations of EWSR1 in the assay buffer (20 mM Bis-Tris-Propane, pH 8) for 30 min at 25 °C. Following the incubation, the nsP2-EWSR1 mix was added to a Nunc 96-well black plate (Thermo Fisher Scientific, 137101) with 25 μM peptide substrate in the assay buffer and the fluorescence was measured at 30 sec intervals using a multimode plate reader SpectraMax i3x at an excitation wavelength of 340 nm and emission wavelength of 490 nm. A substrate control reaction measured the auto-fluorescence generated by just the substrate. Withanone was used at 10 μM concentration as a negative control, and withaferin A at 10 μM concentration as the positive control for the protease activity inhibition assay (22). All enzyme reactions were performed in duplicates and two independent experiments. The average fluorescence of each sample was calculated and plotted in the graph.

### Statistical analysis

The statistical significance of difference in the mean of the multiple data values was determined by unpaired Student’s t-test and indicated in the figures as *, *p*<0.05; **, *p*<0.01; ***, *p*<0.001; ns=not significant.

## RESULTS

### CHIKV genome has alphavirus-conserved genome nucleotide sequences

Three previously reported alphavirus-conserved genomic RNA sequences (8, 11–13) were examined for their conservation in the CHIKV RNA genome. The 40-nucleotide 3NCR40 sequence (nucleotide 11581-11620) is part of the conserved elements and RSEs found upstream in the 3′-NCR. The 48-nucleotide 3NCR48 sequence represents the extreme 3’-end sequence in the CHIKV 3’-NCR (nucleotides 11751-11798). The 51-nucleotide 5NCR51 located from nucleotide 153 to 203 in the CHIKV nsP1-coding region is downstream of the 44-nucleotide putative minus-sense RNA promoter and is critical for efficient virus replication (15, 28, 29). The 50-nucleotide 5NCR-RC50 has sequence complementary to the 5’-NCR sequence and may contain elements required for the plus-sense genomic RNA synthesis (9, 11, 30). Alignment of these CHIKV sequences showed significant sequence conservation across CHIKV-related arthritogenic alphaviruses, such as Ross River virus (RRV), O’nyong-nyong virus (ONNV), Getah virus (GETV), and Barmah Forst virus (BFV) for the 3NCR40, 3NCR48, and 5NCR51 sequences, whereas for the 5NCR-RC50 the nucleotide sequence conservation was mostly patchy and poor (Fig. 1).

**Figure 1.**
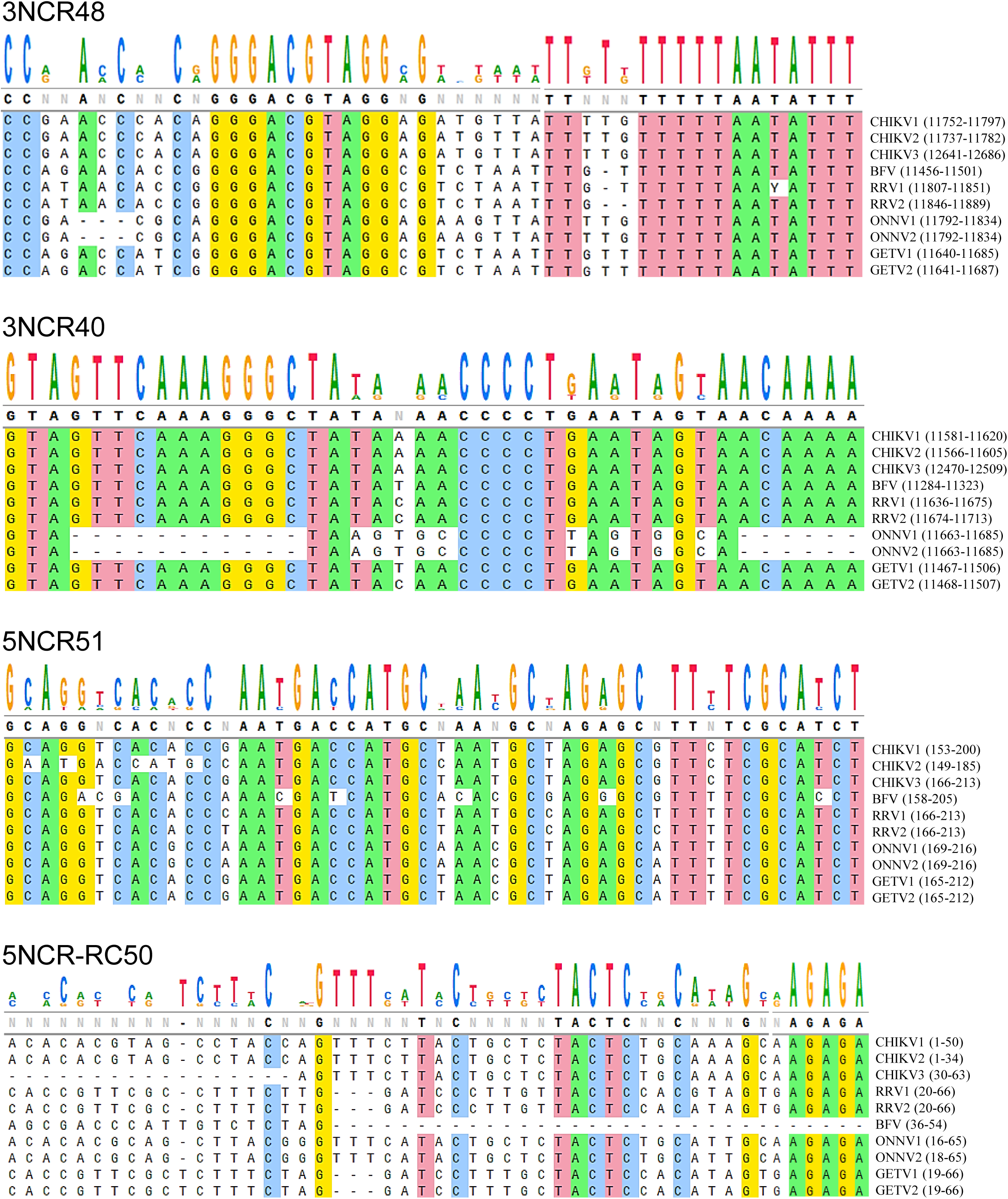
Conserved sequences in the alphavirus genomes. The 3’- and 5’-NCR sequences in CHIKV-related arthritogenic alphaviruses, such as RRV, ONNV, GETV, and BFV were aligned using the Clustal software. The figure shows the alignment to demonstrate sequence conservation in the 3’-NCR around CHIKV sequence represented in 3NCR48 and 3NCR40 sequences, and in the 5’-NCR around CHIKV sequences represented in 5NCR51 and 5NCR-RC50. The GenBank accession numbers of the various sequences used for alignment are as follows: CHIKV1, JF274082.1; CHIKV2, MH349097.1; CHIKV3, EU224269.1; BFV, MK697274.1; RRV1, GQ433359.1; RRV2, NC_075016.1; ONNV1, M20303.1; ONNV2, NC_001512.1; GETV1, LC814433.1; GETV2, MT086508.1. The location of the aligned sequence on respective genomes as nucleotide numbers is shown in the brackets.

### Host proteins bind to the conserved CHIKV RNA sequences

EMSA was used to study the binding of proteins from ERMS, HEK293T, and Vero cells to different CHIKV RNAs. While a number of host proteins from these cells showed binding to 3NCR40, 3NCR48, and 5NCR-RC50, very little host protein binding was seen with 5NCR51 (Fig. 2). Some of the RNA-protein bands diminished or disappeared in the presence of excess polyIC, indicating a non-specific or weak binding. However, several RNA-protein bands were not affected even in the presence of excess polyIC, indicating a high affinity specific binding of the protein with the CHIKV RNAs. Importantly, the binding profile was different with the cell lysates from different cells. The binding profile of the lysate from a given cell line was also different for different RNAs. These data suggest that different host cell proteins may bind to different CHIKV conserved RNA sequences, and different proteins from different cells may bind to a given RNA sequence.

**Figure 2.**
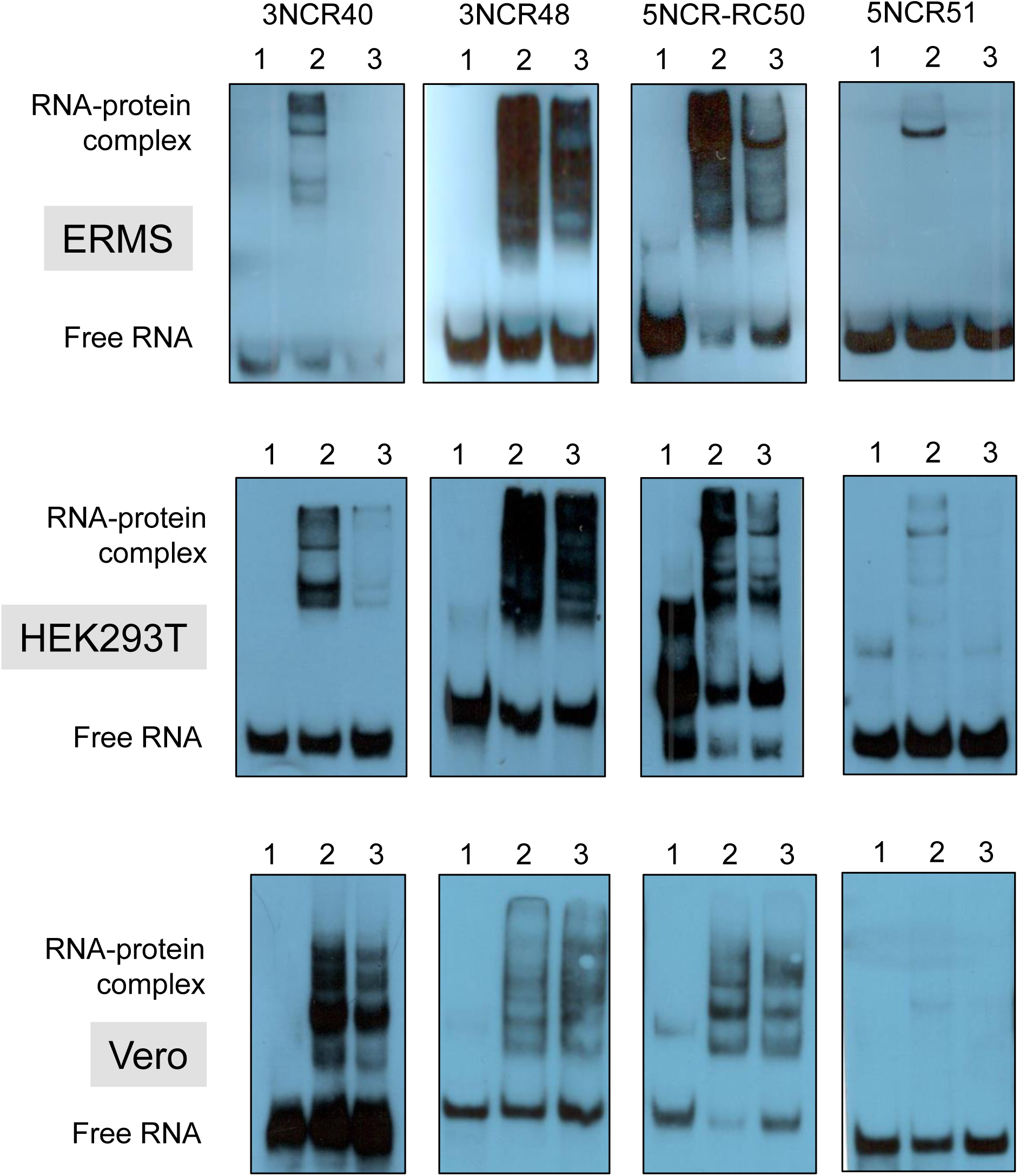
Host proteins bind to the conserved CHIKV RNA sequences. EMSA was used to study the binding of ERMS, HEK293T, and Vero cell proteins to 3NCR40, 3NCR48, 5NCR-RC50, and 5NCR51. Biotin-labelled synthetic RNA (20 ng) was incubated in the binding buffer with 10 μg cell lysate in presence or absence polyIC. The RNA-protein complexes were electrophoresed on a nondenaturing 12% polyacrylamide gel and transferred to the membrane. The RNA and the RNA-protein complexes were visualized using a chemiluminescent nucleic acid detection method. Lane 1, RNA; lane 2, RNA-protein complexes in the absence of polyIC; lane 3, RNA-protein complexes in presence of 10 μg polyIC.

### Identification of the host proteins binding to CHIKV RNA sequences

The host proteins interacting with the biotin-labelled CHIKV RNAs were pulled down from the host cell lysates using streptavidin beads and mass spectrometry was used to identify the proteins. The specific proteins binding to the CHIKV RNAs were identified by eliminating those proteins that bind to the beads or a random RNA. The host proteins identified at least four times from the six experimental replicates are listed in table 1. It can be seen that a diverse set of proteins were binding to different CHIKV RNAs. As was seen in the EMSA, different proteins from different cells were binding to a given RNA sequence, except for CPSF7 that bound 3NCR48 in Vero, ERMS and HEK293T cell lysates, and QKI bound 3NCR40 in both ERMS and HEK293T cell lysates. Notably, only a smaller number of proteins were binding to 5NCR51 RNA. This diversity in the identified proteins establishes the specificity of the RNA-protein binding. The PANTHER analysis was done to understand the molecular function of these proteins binding to the CHIKV RNAs (Table 2). As expected, a majority of the proteins had nucleic acid binding property, in particular RNA.

**Table 1.** Host proteins binding to the CHIKV RNAs.

| **CHIKV RNA** | **Vero cells** | **ERMS cells** | **HEK293T cells** |
| --- | --- | --- | --- |
| 3NCR40 | ROA3 (4/6)<br>QOR (4/6)<br>DAZP1 (4/6) | QKI (6/6) | QKI (6/6)<br>PSPC1 (6/6)<br>HNRDL (6/6)<br>ROA0(6/6) |
| 3NCR48 | CPSF7(6/6)<br>ELAV1 (6/6)<br>PUF60 (6/6)<br>RBMS1 (6/6)<br>ROA0(6/6)<br>SRSF9 (6/6)<br>TIA1 (6/6) | CPSF6 (6/6)<br>CPSF7(6/6) | EWS (6/6)<br>CPSF6 (6/6)<br>CPSF7 (6/6)<br>HNRH3 (6/6) |
| 5NCR51 | IMDH2 (6/6) | SSBP1 (5/6) | CAPR1 (4/6) |
| 5NCR-RC50 | FUS (6/6) | HNRDL (6/6) | EWS (6/6)<br>G3BP1 (6/6) |

The number of times the protein was identified out of the multiple experimental replicates is shown in the brackets.

**Table 2.** PANTHER analysis of the CHIKV RNA binding proteins.

| **GO molecular function** | **Proteins** |
| --- | --- |
| RNA binding | ROA3, QOR, DAZP1, QKI, PSPC1, HNRDL, ROA0, CPSF7, ELAV1, PUF60, RBMS1, SRSF9, TIA1, EWSR1, CPSF6, HNRH3, IMDH2, SSBP1, CAPR1, FUS, HNRDL, G3BP1 |
| mRNA binding | ROA0, CAPR1, TIA1, CPSF6, FUS, ROA3, G3BP1, RBMS1, ELAV1, QKI, QOR, DAZP1, CPSF7 |
| mRNA 3'-NCR binding | ROA0, TIA1, FUS, ROA3, RBMS1, ELAV1, QKI, QOR, DAZP1 |
| nucleic acid binding | ROA0, CAPR1, HNRDL, TIA1, PSPC1, CPSF6, FUS, IMDH2, EWSR1, PUF60, ROA3, G3BP1, RBMS1, ELAV1, SSBP1, SRSF9, QKI, HNRH3, QOR, DAZP1, CPSF7 |
| poly-purine tract binding | HNRDL, TIA1, RBMS1, DAZP1 |
| single-stranded RNA binding | HNRDL, TIA1, RBMS1, DAZP1 |
| poly(A) binding | HNRDL, TIA1, RBMS1 |
| mRNA 3'-NCR AU-rich region binding | ROA0, TIA1, ELAV1 |
| molecular condensate scaffold activity | CAPR1, FUS, G3BP1 |
| poly(G) binding | HNRDL, DAZP1 |

Interestingly EWSR1, detected in all six experimental replicates, was interacting with both the plus- and minus-sense CHIKV RNAs, represented by 3NCR48 and 5NCR-RC50, respectively, in HEK293T cell lysate. We, therefore, further studied the EWSR1 binding to the CHIKV RNAs and its role in CHIKV replication in HEK293T cells.

### EWSR1 shows binding to 3NCR48 and 5NCR-RC50 sequences

Recombinant EWSR1 produced in *E. coli* was used to study its binding to 3NCR48 and 5NCR-RC50 sequences by EMSA. RNA-protein complexes were seen when EWSR1 was incubated with 3NCR48 or 5NCR-RC50 RNAs (Fig. 3). These RNAs bound specifically to EWSR1 as no RNA-protein complex was seen with a random protein BSA. The recombinant EWSR1 used in these assays has the histidine tag. To rule out the role of the histidine tag in RNA binding, EMSA was done using an unrelated, histidine-tagged mycobacterial protein polyphosphate kinase 1 (PPK1), produced in *E. coli*. PPK1 showed no binding to 3NCR48 or 5NCR-RC50. The RNA-protein binding specificity was further established as EWSR1 showed no binding with Random48, a 48-nucleotide non-specific RNA with randomized sequence derived from 3NCR48.

**Figure 3.**
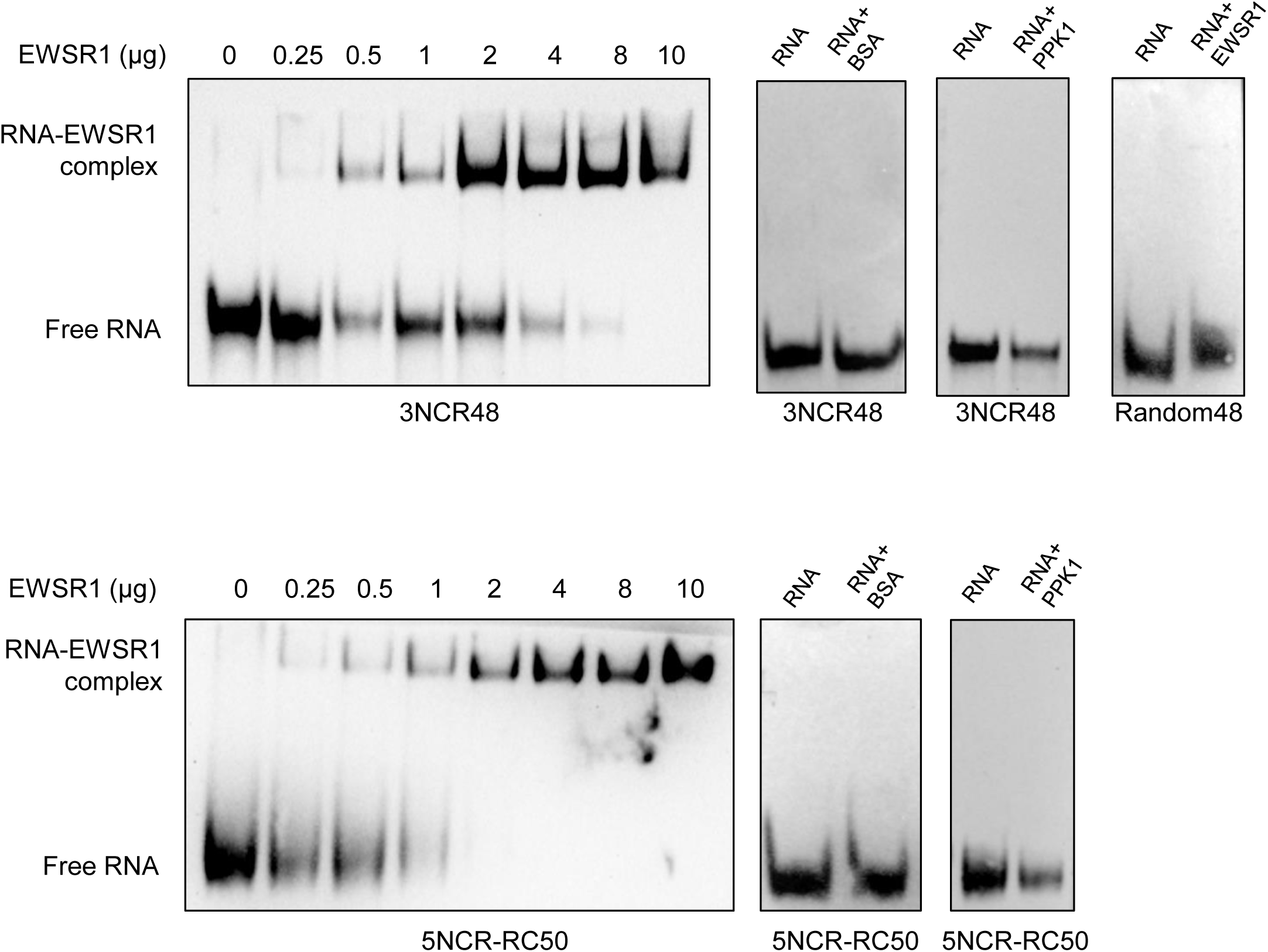
EWSR1 binding with CHIKV RNAs. EMSA was used to study the binding of EWSR1 to 3NCR48 and 5NCR-RC50. Biotin-labelled synthetic RNA (10 ng) was incubated in the binding buffer with indicated amounts of purified protein in presence of 10 μg polyIC. The RNA-protein complexes were electrophoresed on a nondenaturing 12% polyacrylamide gel and transferred to the membrane. The RNA and the RNA-protein complexes were visualized using a chemiluminescent nucleic acid detection method. BSA and PPK1 (10 μg each) and Random48 (10 ng) were used as the non-specific protein and RNA controls, respectively.

The RNA-protein binding was further studied using the label-free biolayer interferometry (BLI) method where biotinylated CHIKV RNAs were immobilized on a streptavidin-coated sensor, and then exposed to different concentrations of EWSR1 (Fig. 4). The protein was binding to both 3NCR48 and 5NCR-RC50 RNAs with high affinity. The equilibrium dissociation constant (*K*_d_) was 91.4±0.37 nM and 80.2±0.39 nM for the EWSR1 binding to 3NCR48 and 5NCR-RC50 RNAs, respectively. The nanomolar *K*_d_ values suggest that these RNA-protein interactions may be physiologically plausible. In these experiments also, 3NCR48 and 5NCR-RC50 RNAs showed insignificant or no binding with a random control protein BSA, and EWSR1 did not bind the control RNA, Random48. These data were consistent with the EMSA observations above.

**Figure 4.**
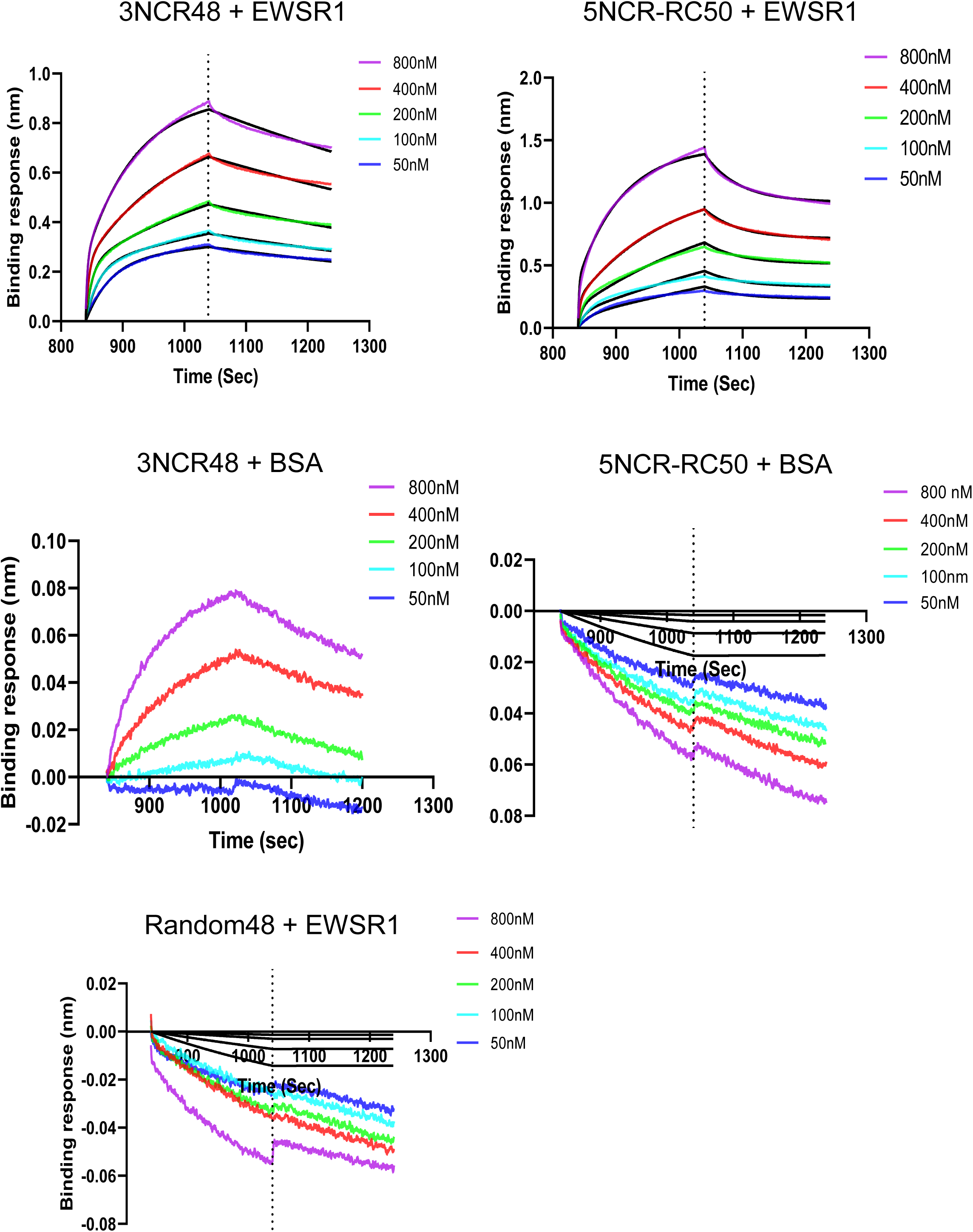
EWSR1 binding to the CHIKV RNAs. Biotinylated CHIKV RNAs 3NCR48 and 5NCR-RC50 (50 ng) were immobilized on the streptavidin-coated biosensor tips and exposed to the increasing concentrations of EWSR1. Representative BLI binding curves showing association and dissociation are presented. BSA and Random48 were used as the non-specific protein and RNA controls, respectively.

### EWSR1 binds CHIKV RNA in the virus-infected cells

RNA immunoprecipitation was employed to examine whether EWSR1 indeed bound CHIKV RNA during the virus infection. Cell lysate from CHIKV-infected cells was incubated with EWSR1 antibody or the matching isotype IgG control and the antibody-protein complex pulled down. The complex was used to extract RNA and qRT-PCR was done to determine the relative level of CHIKV RNA (Fig. 5). A significantly increased (*p*<0.001) amount CHIKV RNA (∼15-fold higher) was detected in the complex pulled-down by EWSR1 antibody, suggesting that EWSR1 was associated with CHIKV RNA in the virus-infected cells.

**Figure 5.**
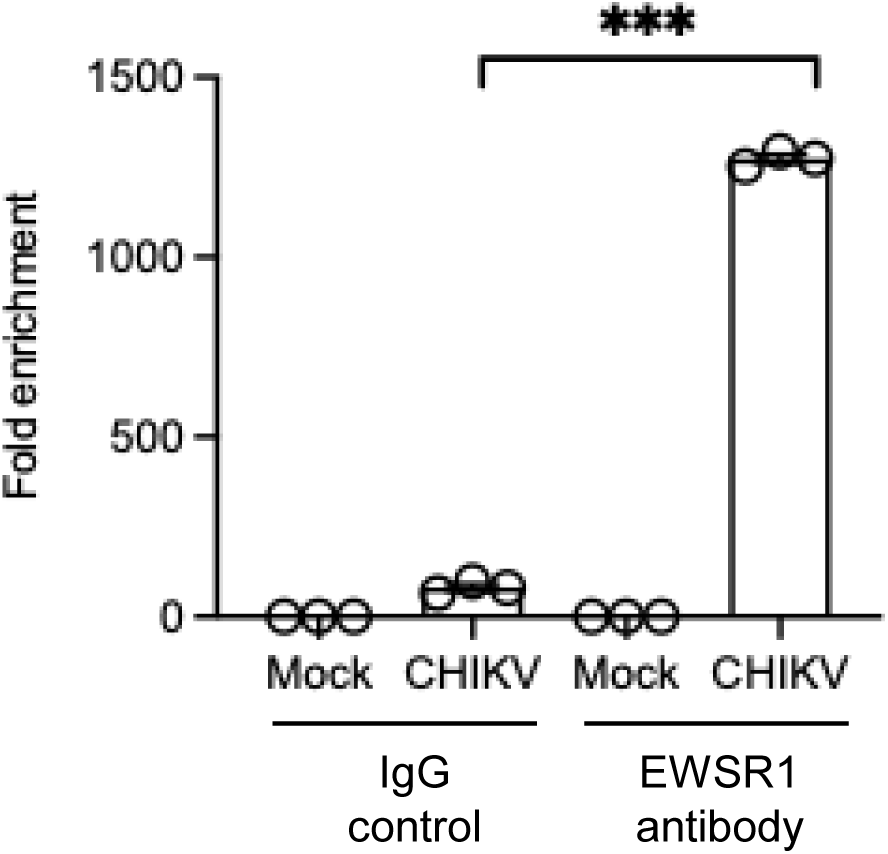
EWSR1 binds CHIKV RNA in the virus-infected cells. HEK293T cells were infected with CHIKV (MOI 0.5) or were mock infected, and the cell lysates were made at 12 h pi. The cell lysates from the equal number of cells were subjected to immunoprecipitation using anti-EWSR1 antibody or an IgG isotype control. The co-immunoprecipitated RNA was extracted and analyzed by qRT-PCR using CHIKV specific primers. Data are presented as relative RNA enrichment compared to that in the mock-infected cell lysate treated with the control IgG.

### CHIKV infection modulates EWSR1 expression

The effect of CHIKV infection on EWSR1 expression was studied by determining the EWSR1 RNA levels by qRT-PCR and protein levels by western blotting (Fig. 6A). As the infection progressed, the EWSR1 RNA and protein levels decreased. EWSR1 is located in both the nucleus and cytoplasm (31). However, during the CHIKV infection, the EWSR1 levels in the nucleus were reduced and its cytoplasmic levels increased (Fig. 6B). These data showed that CHIKV infection downregulated the expression of EWSR1 RNA and protein, and resulted in the EWSR1 protein accumulation in the cytoplasm.

**Figure 6.**
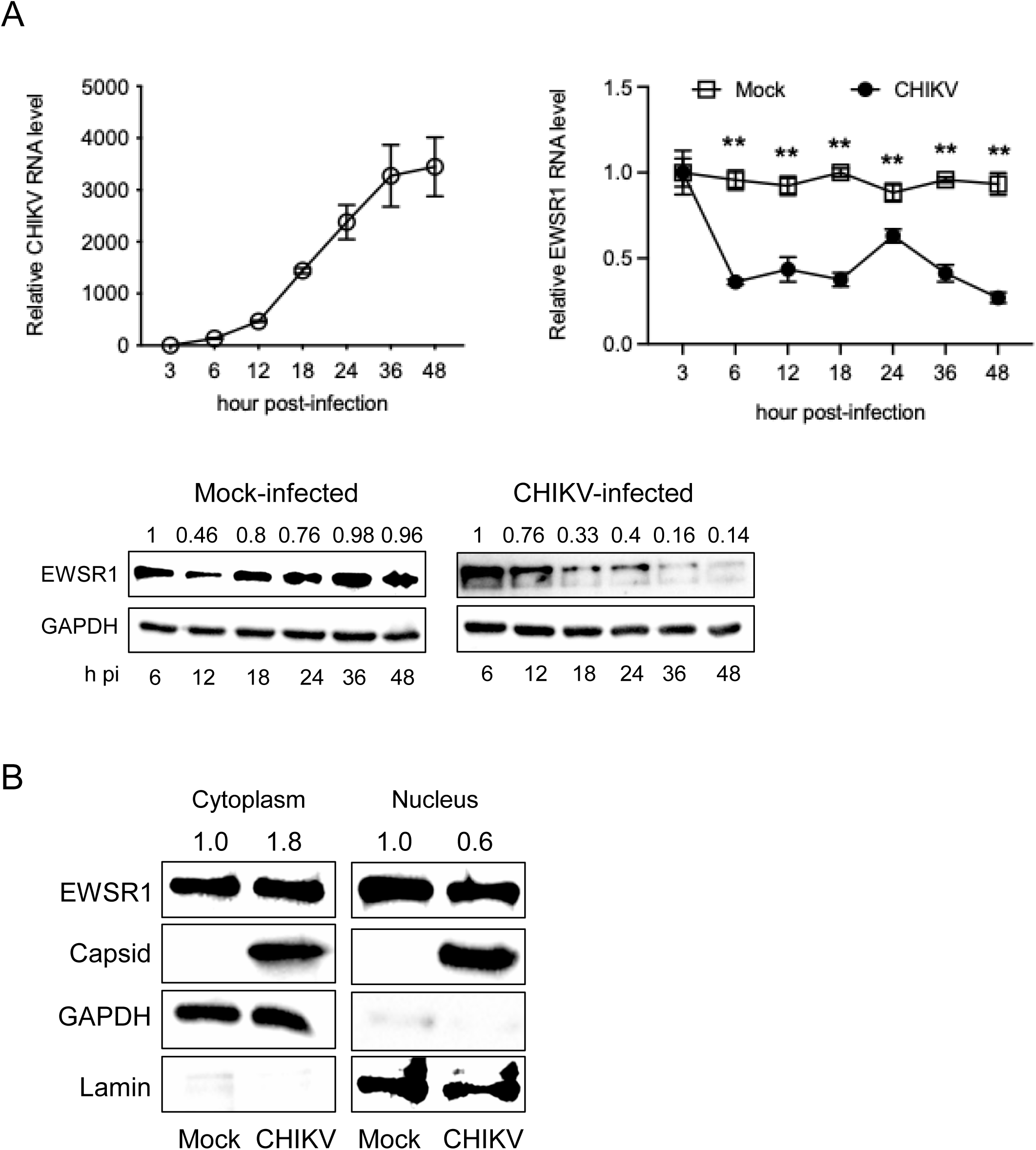
CHIKV infection modulates EWSR1 expression. (A) HEK293T cells were mock-infected or infected with CHIKV (MOI 0.5). The cells and culture supernatants were harvested at different times pi. CHIKV titers are shown in the top left panel. The cell lysates were used to extract the total RNA and the EWSR1 RNA levels in the mock- and virus-infected cells determined by qRT PCR. GAPDH was used as the internal control for normalization. EWSR1 RNA levels relative to that seen at 3 h pi are presented in the top right panel. The cell lysates were western blotted to determine the EWSR1 protein levels in the mock- and virus-infected cells. GAPDH levels were used for normalization. The intensity of EWSR1 bands relative to GAPDH are shown at the top where the EWSR1/GAPDH ratio at 6 h pi was taken as one. (B) HEK293T cells were mock-infected or infected with CHIKV (MOI 0.5). The cells were harvested at 12 h pi and the lysates were separated into the nuclear and cytoplasmic fractions. GAPDH and Lamin were blotted to establish the purity of the cytoplasmic and nuclear fractions, respectively. The relative levels of EWSR1 protein compared to GAPDH (in cytoplasm) and Lamin (in nucleus) are shown at the top. The relative value in the mock-infected cells was taken as 1.

### EWSR1 has an antiviral role in CHIKV replication

The siRNA-mediated knockdown of EWSR1 was used to study its role in CHIKV replication. HEK293T cells were treated with siEWSR1 or siNT (non-targeting control) for 48h. This resulted in ∼60% reduction in the EWSR1 mRNA levels and ∼75% reduction in the protein levels (Fig. 7A). At this point, the cells were infected with CHIKV, and the cells and supernatant were harvested at different times post-infection (pi) to determine the viral RNA and titer, respectively. While a 35% enhanced CHIKV RNA levels were seen at 12 h pi, the virus titer was unaffected in the siEWSR1-treated cells at this time point (Fig. 7B). In another experiment, CHIKV titers were measured at later time points of 18 and 24 h pi (Fig. 7C). Here also, viral titers were not affected at 12 h pi although these were significantly lower (*p*<0.05) by 30-40% at 18 and 24 h pi in the siEWSR1-treated cells than the control.

**Figure 7.**
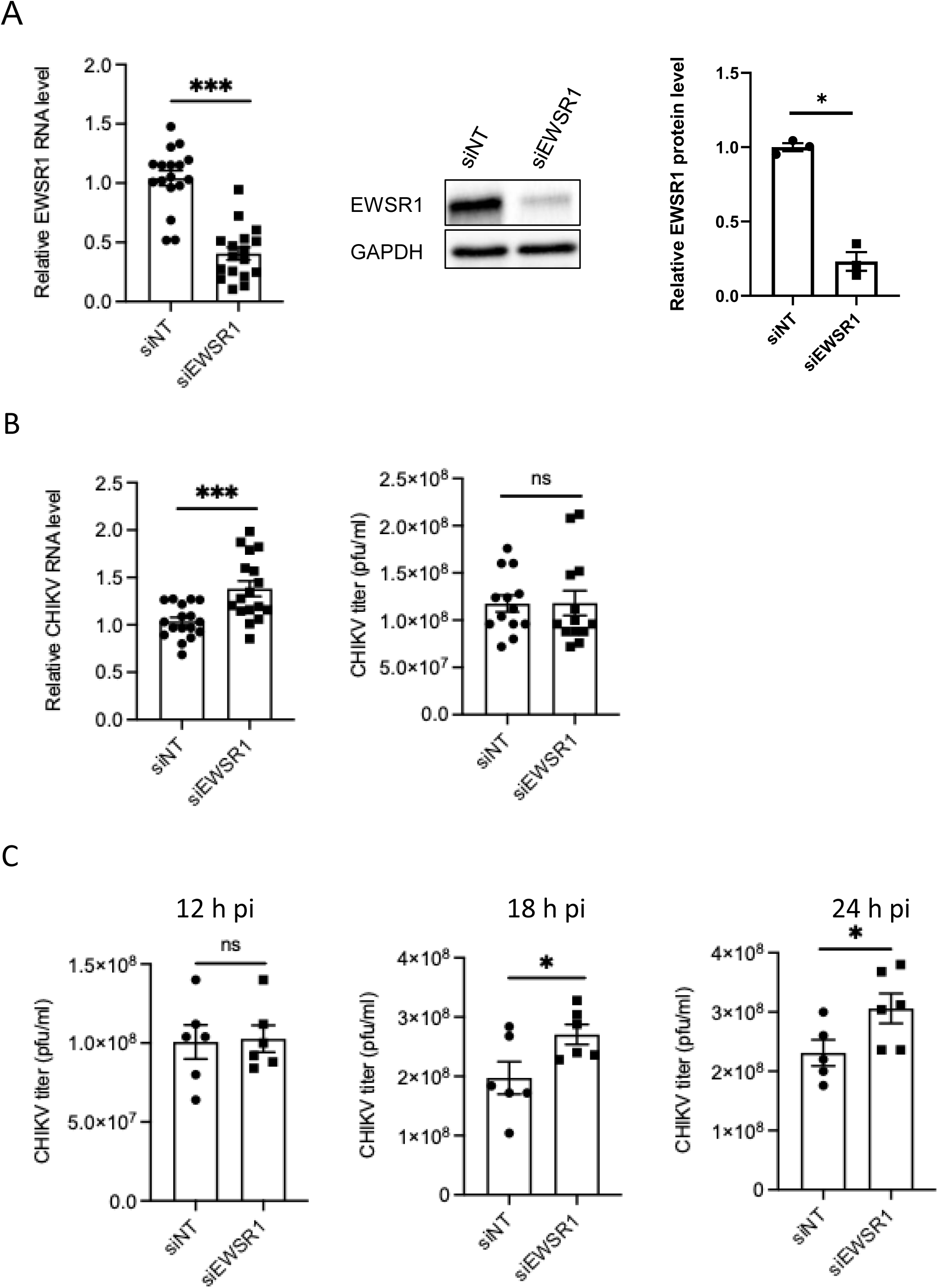
CHIKV replication in the siRNA-mediated EWSR1 downregulated cells. (A) HEK293T cells were transfected with siEWSR1 or a control non-targeting siNT and harvested 48 h later to determine the relative EWSR1 RNA and protein levels by qRT PCR and western blotting, respectively. The left panel shows the relative EWSR1 RNA levels. The right panel shows the relative EWSR1 protein levels based on the band intensity in the western blots. A representative western blot is shown in the middle. (B) HEK293T cells were transfected with siEWSR1 or siNT and 48 h later infected with CHIKV (MOI 0.5). The cells and culture supernatants were harvested 12 h later. The CHIKV RNA levels in siEWSR1-treated cells relative to the siNT-treated cells are shown in the left panel. CHIKV titers are shown in the right panel. (C) HEK293T cells were transfected with siEWSR1 or siNT and 48 h later infected with CHIKV (MOI 0.5). The culture supernatants were harvested at indicated time points. The CHIKV titers in the siNT- and siEWSR1-treated cells are shown.

We also studied the effect of EWSR1 overexpression on CHIKV replication. HEK293T cells were transfected with a plasmid ectopically expressing EWSR1 which resulted in 120% increase in the EWSR1 RNA levels and 95% enhanced EWSR1 protein levels at 24 h post-transfection (Fig. 8A). At this point, cells were infected with CHIKV, and 12 h pi the samples were harvested to determine the CHIKV RNA and the virus titers. A significant decrease in CHIKV RNA by 35% (*p*<0.001) and titers by 40% (*p*<0.01) was seen in EWSR1-overexpressing cells (Fig. 8B).

**Figure 8.**
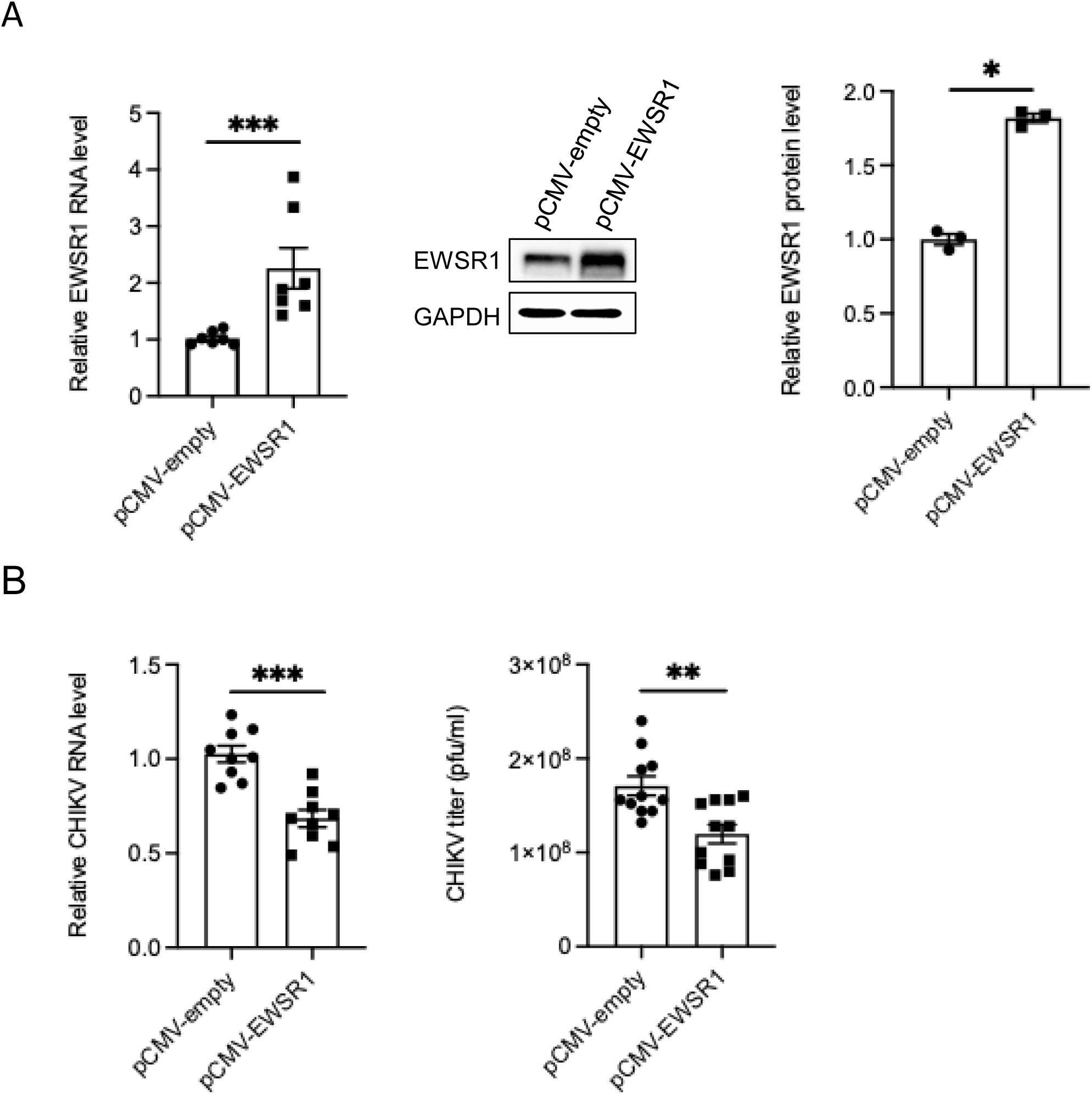
CHIKV replication in the ectopically-EWSR1 expressing cells. (A) HEK293T cells were transfected with the empty vector pCMV-empty or EWSR1-expressing pCMV-EWSR1 and harvested 24 h later to determine the relative EWSR1 RNA and protein levels by qRT PCR and western blotting, respectively. The left panel shows the relative EWSR1 RNA levels. The right panel shows the relative EWSR1 protein levels A representative western blot is shown in the middle. (B) HEK293T cells were transfected with pCMV-empty or pCMV-EWSR1 and 24 h later infected with CHIKV (MOI 0.5). The cells and culture supernatants were harvested 12 h later. The CHIKV RNA levels in pCMV-EWSR1-transfected cells relative to the pCMV-empty transfected cells are shown in the left panel. CHIKV titers are shown in the right panel.

To validate these findings and to rule out any off-target effects of siEWSR1, an EWSR1 knockout (KO) of HAP1 cells was made. The KO cells were infected with CHIKV, and 12 h pi the cells and supernatant were collected to determine the viral RNA levels and titers. The CHIKV RNA levels and titers were significantly increased by 25% (*p*<0.05) and 115% (*p*<0.01), respectively, in the KO cells compared to the wild-type (WT) control (Fig. 9A). We also rescued the EWSR1 expression in the EWSR1 KO HAP1 cells by transfecting them with a plasmid ectopically expressing EWSR1. In the EWSR1-rescued KO cells (KOR) the CHIKV RNA levels were reduced by 90% (p<0.01) and virus titers by 35% (*p*<0.05), compared to the EWSR1 KO cells (Fig. 9B). Together, these data demonstrate an antiviral role of EWSR1 in CHIKV replication.

**Figure 9.**
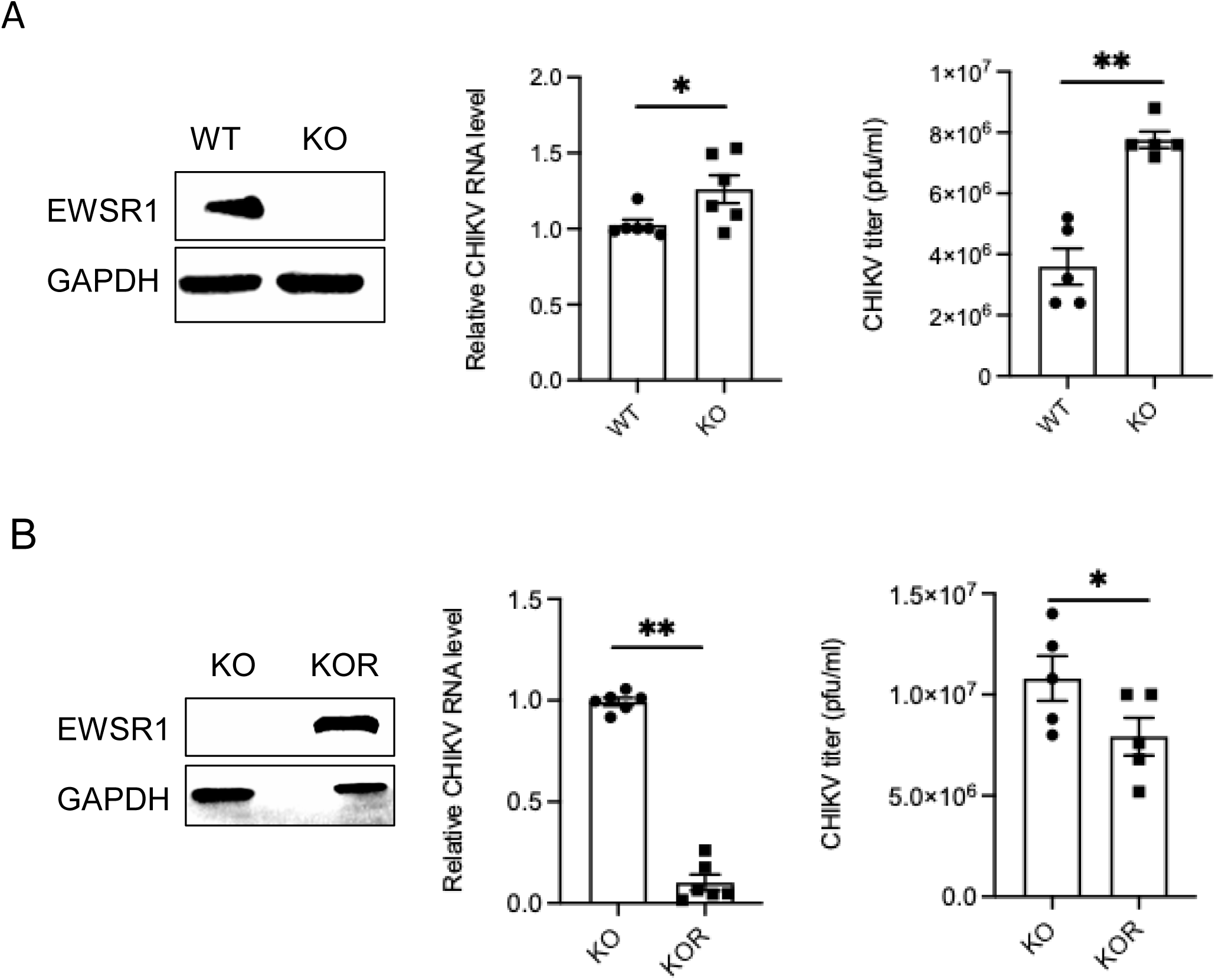
CHIKV replication in the EWSR1 knockout HAP1 cells. (A) Wild type (WT) and EWSR1 knockout (KO) HAP1 cells were cultured and western blotted to establish the absence of EWSR1 protein in the KO cells (left panel). The WT and KO HAP1 cells were infected with CHIKV (MOI 0.5) and 12 h later CHIKV RNA levels in the KO cells relative to that in the WT cells were determined by qRT PCR (middle panel). The CHIKV titers in the WT and KO cells are presented in the right panel. (B) EWSR1 KO HAP1 cells were transfected with a EWSR1 expressing pCMV-EWSR1. The EWSR1 KO and EWSR1-rescued cells (KOR) were western blotted 24 h later using the EWSR1 antibody to establish the presence of the ectopically-expressed EWSR1 (left panel). The KO and KOR cells were infected with CHIKV (MOI 0.5) and the cells and culture supernatants were harvested 12 h later. The viral RNA levels in the KOR cells relative to the KO cells are shown (middle panel). The viral titers are shown in the right panel.

### EWSR1 reduces both plus- and minus-sense CHIKV RNA levels in the virus-infected cells

EWSR1 can bind both plus- and minus-sense CHIKV RNAs. How this binding might affect the synthesis of these RNAs during the virus replication was studied in HAP1 KO and WT cells (Fig. 10A). In the KO cells, both minus- and plus-sense CHIKV RNA copy numbers were significantly higher than those in the WT cells (*p*<0.05 or 0.01 at different time points). Ectopic expression of EWSR1 in the KO cells (KOR) significantly reduced both the minus-and plus-sense CHIKV RNA copy numbers (*p*<0.05 or 0.01 at different time points) (Fig. 10B). This was further validated by ectopic expression of EWSR1 in HEK293T cells where cells over-expressing EWSR1 showed significantly reduced copy numbers both for the minus- and plus-sense CHIKV RNAs (*p*<0.05, 0.01, or 0.001 at different time points). (Fig. 10C). Since EWSR1 binds sequences within the plus- and minus-sense CHIKV RNAs, this might have a role in the reduced viral RNA synthesis resulting in the lowering of CHIKV titers.

**Figure 10.**
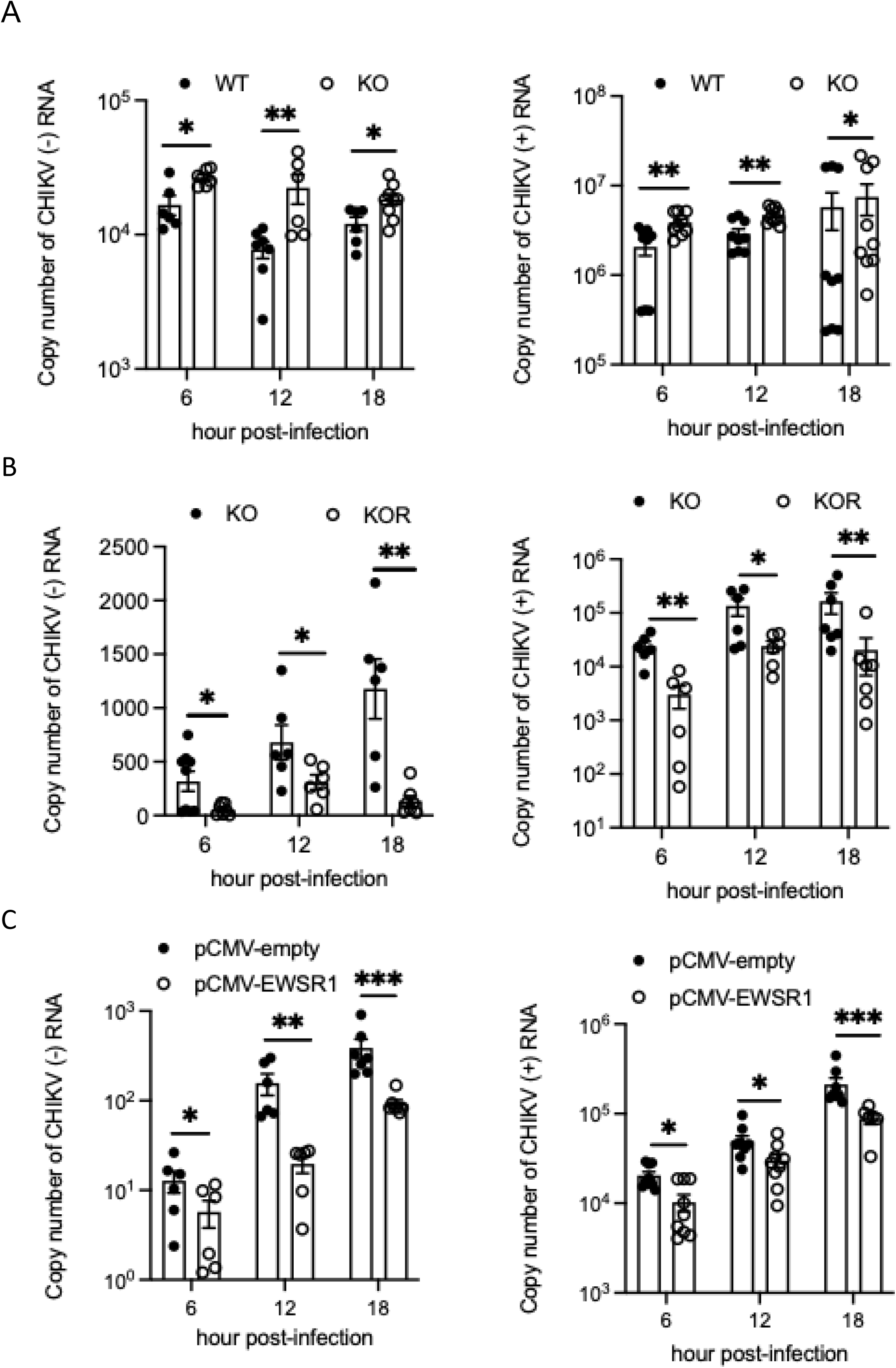
CHIKV plus- and minus-sense RNA copy numbers in the virus-infected cells. (A) The EWSR1 WT and KO HAP1 cells were infected with CHIKV (MOI 0.5) and the cells were harvested at different times pi to isolate the total RNA. qRT PCR was done to determine the CHIKV plus- and minus-sense RNA copy numbers. (B) The EWSR1 KO and the KOR HAP1 cells were infected with CHIKV (MOI 0.5) and the cells were harvested at different times pi to isolate the total RNA. qRT PCR was done to determine the CHIKV plus-and minus-sense RNA copy numbers. (C) HEK293T cells were transfected with the empty vector pCMV-empty or EWSR1-expressing pCMV-EWSR1, and 24 h later infected with CHIKV (MOI 0.5). The cells were harvested at different times pi to isolate the total RNA. qRT PCR was done to determine the CHIKV plus- and minus-sense RNA copy numbers.

### CHIKV nsP2 binds 3NCR48 sequence

EWSR1 binds the 3NCR48 and 5NCR-RC50 sequences which are at the extreme 3’-end of the CHIKV plus-sense genomic RNA and minus-sense replication-intermediate RNA, respectively. These ends interact with alphavirus non-structural proteins to initiate the RNA synthesis (11). In fact, nucleotide sequence GUUUUUAAUAUUUC, which is part of the 3NCR48 sequence, has been shown to bind the CHIKV nsP2 helicase (32). We therefore sought to examine if CHIKV nsP2 binds the 3NCR48 RNA. Indeed, EMSA showed CHIKV nsP2 binding to the 3NCR48 sequence (Fig. 11A). We then studied the binding of 3NCR48 RNA with CHIKV nsP2 by BLI which validated the CHIKV nsP2 binding with the 3NCR48 RNA (Fig. 11B). The equilibrium dissociation constant (*K*_d_) for the CHIKV nsP2 binding with 3NCR48 RNA was 92.9±1.6 nM.

**Figure 11.**
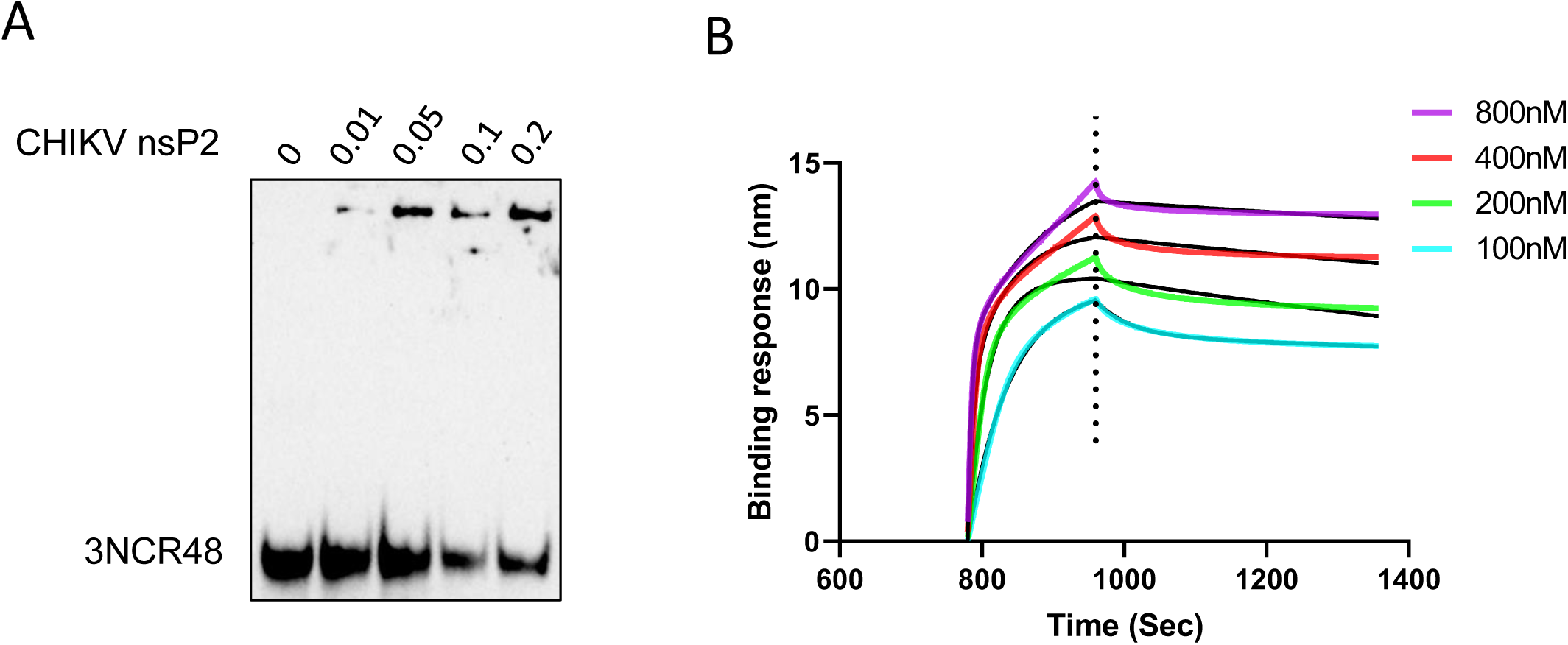
CHIKV nsP2 binds the 3NCR48 RNA. (A) EMSA was used to study the binding of CHIKV nsP2 with 3NCR48 RNA. Biotin-labelled synthetic RNA (20 ng) was incubated in the binding buffer with different amounts of CHIKV nsP2 protein in presence of polyIC. The RNA-protein complexes were electrophoresed on a nondenaturing 12% polyacrylamide gel and transferred to the membrane. The RNA and the RNA-protein complexes were visualized using a chemiluminescent nucleic acid detection method. (B) Biotinylated 3NCR48 RNA (50 ng) was immobilized on the streptavidin-coated biosensor tips and exposed to the increasing concentrations of CHIKV nsP2. Representative BLI binding curves showing association and dissociation are presented.

### EWSR1 binds to CHIK nsP2

In a recent study, EWSR1 was shown to interact with the SARS-CoV2 helicase protein (33). We, therefore, examined if EWSR1 was interacting with CHIKV nsP2 protein, which has helicase activity. From the CHIKV-infected cell lysate, we immunoprecipitated EWSR1-interacting protein/s, if any, using the EWSR1 antibody. Interestingly, we detected CHIKV nsP2 protein in the immunoprecipitate (Fig. 12A), suggesting that EWSR1 may be interacting with CHIKV nsP2 during the virus infection.

**Figure 12.**
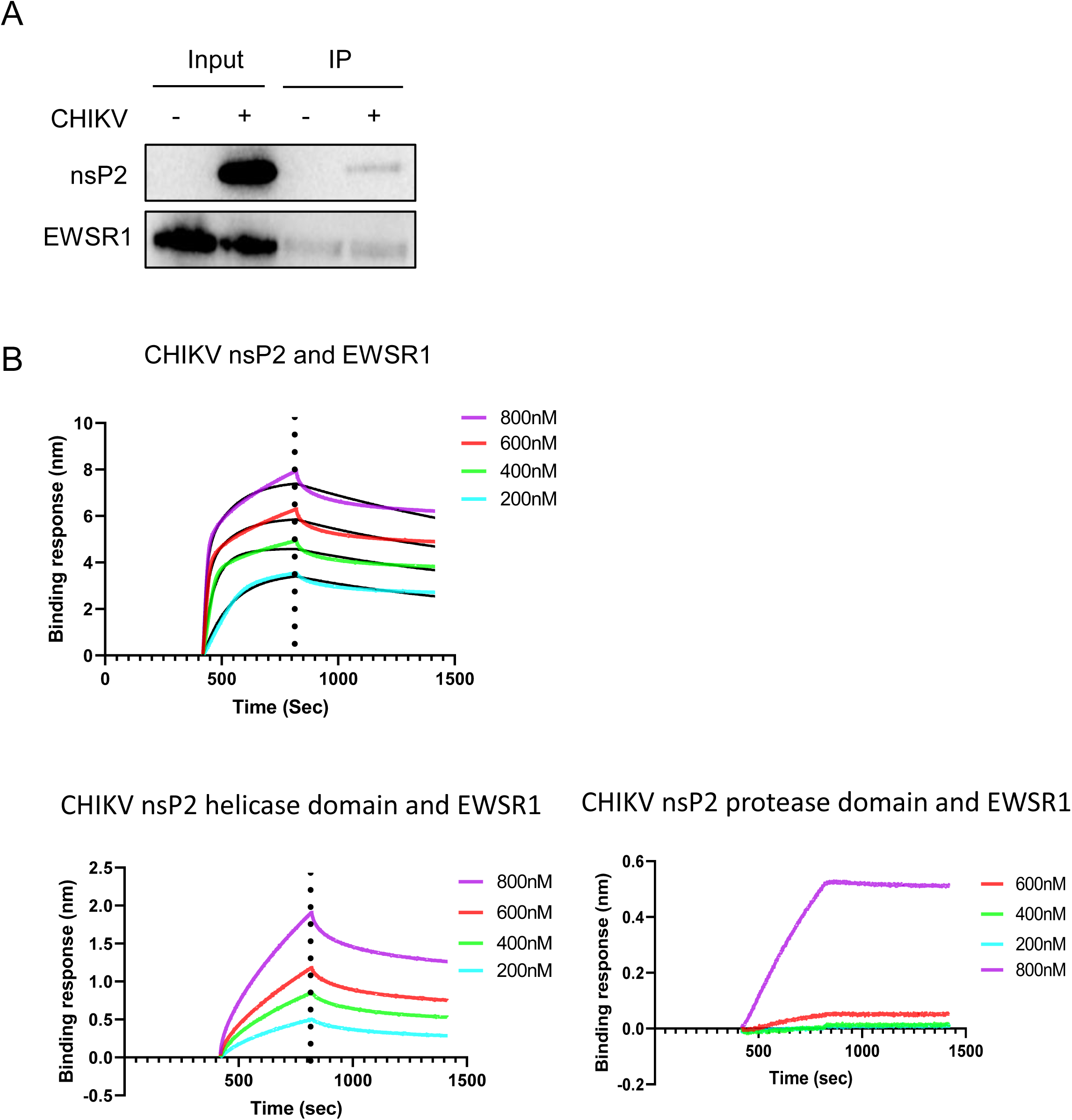
EWSR1 binds to CHIK nsP2. (A) HEK293T cells were mock-infected (-) or infected (+) with CHIKV (MOI 0.5) and 12 h later cell lysates (input) were prepared which were subjected to immunoprecipitation using the EWSR1 antibody. The immunoprecipitate was western blotted with anti-EWSR1, and anti-CHIKV nsP2 antibodies. (B) His-tagged EWSR1 (10 µg) was immobilized on Ni-NTA biosensor tips and exposed to increasing concentrations of CHIKV nsP2 protein, or its helicase and protease domain proteins. Representative BLI binding curves showing association and dissociation of the proteins at different nsP2 concentrations are presented.

BLI method was employed to validate the binding of EWSR1 with CHIKV nsP2 (Fig. 12B). Recombinant histidine-tagged EWSR1 made in *E. coli* was immobilized on a Ni-NTA coated biosensor, and then exposed to different concentrations of CHIKV nsP2 protein purified from the recombinant *E. coli*. The BLI sensorgram demonstrated the binding of EWSR1 with CHIKV nsP2; the equilibrium dissociation constant (*K*_d_) was 716.2±1.76 nM. CHIKV nsP2 is a multifunctional protein having distinct helicase and protease domains (32). The BLI method showed EWSR1 binding with the nsP2 helicase domain. However, no EWSR1 binding was seen with the nsP2 protease domain (Fig. 12B).

### EWSR1 inhibits CHIKV nsP2 helicase activity but not the protease activity

To examine if the EWSR1 binding to CHIKV nsP2 affected its helicase activity, the nsP2 helicase assay was done in the presence of the increasing amounts of EWSR1 (Fig. 13A). As expected, CHIKV nsP2 protein was able to unwind the dsRNA substrate producing ssRNA. The helicase activity of CHIKV nsP2 was inhibited by EWSR1 in a concentration-dependent manner. While partial inhibition of the helicase activity was seen at the lower concentrations, a near complete inhibition was seen in the presence of 1 μM EWSR1 protein.

**Figure 13.**
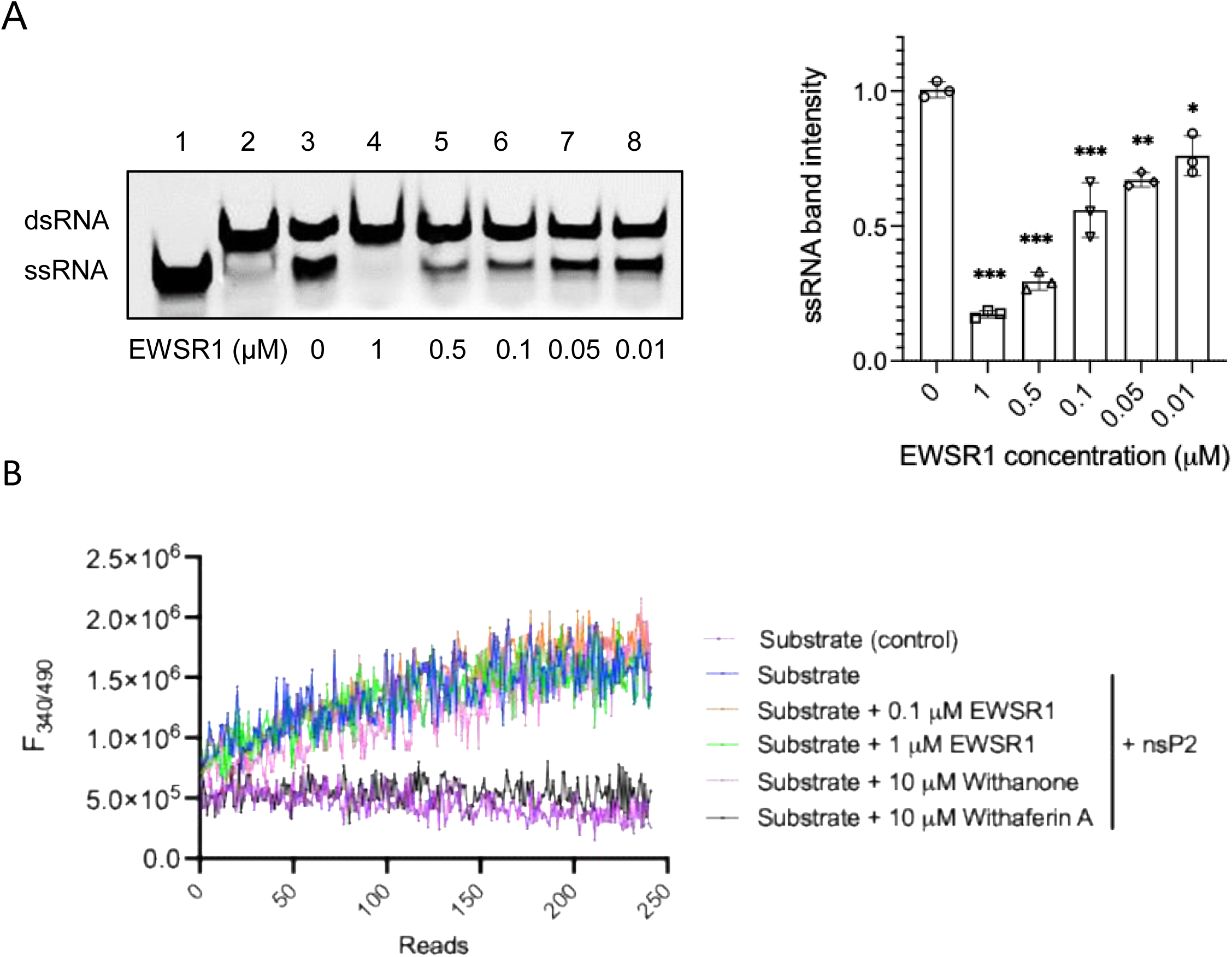
EWSR1 inhibits CHIKV nsP2 helicase activity. (A) The RNA helicase activity of CHIKV nsP2 in presence of increasing EWSR1 concentrations was studied using Alexa488-dsRNA28/16 substrate. The representative gel picture of an experiment is shown in the left panel. Lane 1 has just the dsRNA substrate, lane 2 has the substrate denatured by heat, thus producing ssRNA, lane 3 shows the helicase activity of CHIKV nsP2 in the absence of EWSR1 producing ssRNA, lanes 4-8 show the nsP2 helicase activity inhibition in the presence of the indicated EWSR1 concentrations. (B) A FRET-based protease assay was used to study the nsP2 protease activity. The real-time profile of the proteolytic assay is presented using 25 μM fluorogenic peptide substrate, 1 μM nsP2 protein, with different concentrations of EWSR1. Withaferin A, a known inhibitor of nsP2 protease, and withanone, known to not inhibit the nsP2 protease activity, were used as the controls. The fluorescence was monitored every 30 sec. The background fluorescence of the substrate peptide was monitored without the enzyme and shown as control (substrate).

A FRET-based assay has been established and validated in our lab to study CHIKV nsP2 protease activity (22). EWSR1 had no effect on the nsP2 protease activity when used at 1 μM concentration (Fig. 13B). Here, withaferin A that has been shown to inhibit CHIKV nsP2 protease activity was used as a positive control and withanone as a negative control for the protease activity inhibition, and both controls worked as reported previously (22).

### Role of EWSR1 in the replication of other alphaviruses

Clustal Omega was used to align CHIKV 3NCR48 sequence with the 3’-NCR sequence of SINV and RRV, and LALIGN was used to calculate the sequence conservation (Fig. 14A). CHIKV 3NCR48 sequence is conserved in RRV (82.6% similarity) whereas only poor conservation (51.3% similarity) of this sequence was seen in SINV. In fact, a large insertion of 25 nucleotides was observed in the corresponding SINV sequence when compared with the CHIKV 3NCR48 sequence. We, therefore, chose these two alphaviruses to study the role of EWSR1 interaction with the 3NCR48 sequence in the virus replication.

**Figure 14.**
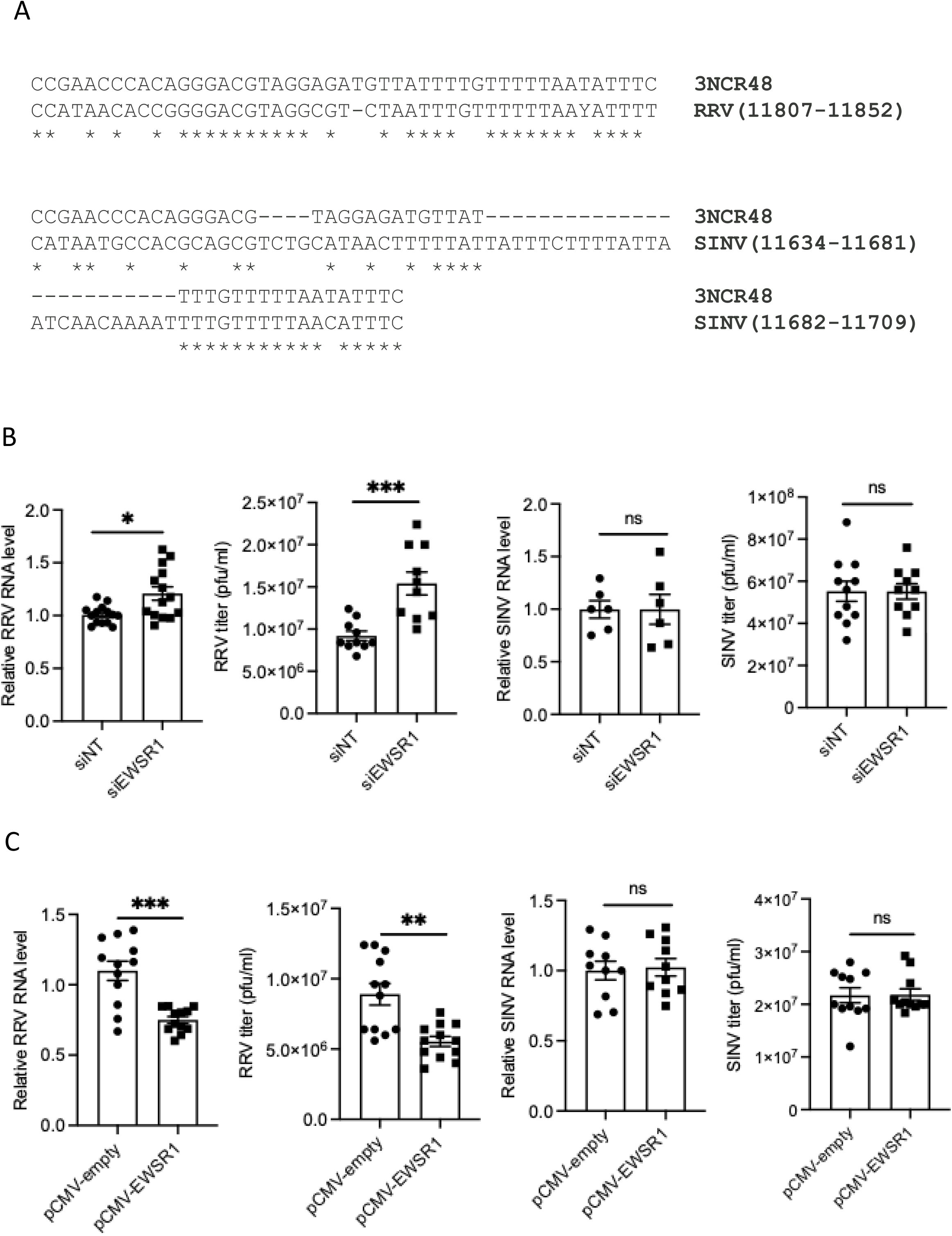
Role of EWSR1 in the replication of RRV and SINV. (A) CHIKV 3NCR48 sequence was aligned with the RRV (GenBank ID GQ433359.1) or SINV (GenBank ID LC869668.1) RNA using Clustal Omega. Conserved nucleotides are identified with an asterisk underneath. The position of the aligned sequence on the RRV or SINV genome is identified as the nucleotide position in the brackets. (B) HEK293T cells were transfected with siEWSR1 or a non-targeting siNT and 24 h later infected with RRV or SINV (MOI 0.5). The cells and culture supernatants were harvested 12 h later. The viral RNA levels in siEWSR1-treated cells relative to the siNT-treated cells are shown. The viral titers are also shown. (C) HEK293T cells were transfected with pCMV-empty or pCMV-EWSR1 and 24 h later infected with RRV or SINV (MOI 0.5). The cells and culture supernatants were harvested 12 h later. The CHIKV RNA levels in pCMV-EWSR1-transfected cells relative to the pCMV-empty transfected cells are shown. The viral titers are also shown.

siRNA-mediated EWSR1-depleted HEK293T cells were infected with RRV or SINV to understand the role of EWSR1 in the replication of other alphaviruses (Fig. 14A). The viral RNA levels and titers were significantly higher (*p*<0.05 and *p*<0.001, respectively) in the siEWSR1-treated RRV-infected cells compared to the siNT-treated RRV-infected control cells. However, SINV RNA levels and virus titers remained unaffected in these cells. Replication of RRV and SINV was also studied in the EWSR1-overexpressing cells (Fig. 14B). While RRV RNA levels and titers were significantly lower (*p*<0.001 and *p*<0.01, respectively) in the cells ectopically overexpressing EWSR1, no effect on SINV RNA and titers was seen. These data, thus, demonstrate the importance of EWSR1 interaction with the 3NCR48 sequence to control the virus replication.

## DISCUSSION

NCRs in the alphavirus genomes play an essential role in the viral replication process. The 5’- and 3’-NCRs contain critical structural and sequence elements that regulate both genome replication and translation (8). Host proteins specifically binding to these sequences, therefore, are likely to have a role in virus replication. Indeed, several host proteins have been shown to bind the CHIKV NCRs and play significant roles in viral replication. Musashi RNA Binding Proteins (MSI-1 and MSI-2) specifically bind to the 5′-NCR of CHIKV RNA. Depletion of these proteins blocks viral replication, showing a proviral role for them (29). Host cellular protein HuR binds alphavirus 3′-NCRs, including CHIKV, stabilizing viral RNA and preventing degradation, thus facilitating virus replication (18). Heterogeneous nuclear ribonucleoprotein K (hnRNP K) is an RNA-binding protein that positively regulates CHIKV replication by modulating the subgenomic RNA (34).

We have identified several host proteins that bind specifically to the synthetic RNAs 3NCR48, 3NCR40, 5NCR51 and 5NCR-RC50 representing sequences located within the CHIKV NCRs or in the immediate vicinity important for RNA replication. Conceivably, the functional annotation of these proteins showed that a majority of these proteins could bind nucleic acid, in particularly RNA. Importantly, different proteins bound to different RNAs establishing the specificity of the RNA-protein binding. Interestingly, one protein, EWSR1, bound both the plus-sense (3NCR48 sequence) and minus-sense (5NCR-RC50 sequence) CHIKV RNAs in HEK293T cells, and therefore, we studied this binding in further details.

The 3NCR48 sequence represents the 48-nucleotide conserved sequence within the alphavirus 3’-NCR. It is located immediately upstream of the poly(A) tail, and functions as a *cis*-acting element that promotes efficient initiation of minus-strand RNA synthesis and supports optimal genome replication and stability (11, 35). Mutagenesis and infectious clone studies show that deletions or alterations in this 48-nucleotide conserved sequence markedly reduce minus-strand synthesis, plaque size, progeny production, and compromise infectivity in both vertebrate and invertebrate hosts, underscoring its essential, evolutionarily conserved role in the alphavirus life cycle (36). The 50-nucleotide 5NCR-RC50 sequence, which is complementary to the 5′-NCR of the virus genome, forms part of the core promoter on the negative-strand template that directs initiation of the plus-sense genomic RNA synthesis, thereby ensuring efficient genome replication during the alphavirus life cycle (37, 38). Mutational analyses show that this region acts as a replication enhancer whose sequence and predicted AU-rich unpaired structure are critical for the positive-strand synthesis in both mosquito and vertebrate cells during the alphavirus replication (39). EWSR1, the host protein binding to the 3NCR48 and 5NCR-RC50 sequences representing the genome sequence elements critical for alphavirus replication, is therefore, likely to affect the virus life cycle.

EWSR1 (Ewing Sarcoma Breakpoint Region 1 / EWS RNA Binding Protein 1) is a ubiquitously expressed multifunctional protein. EWSR1 participates in transcriptional co-regulation and RNA-binding activities, influencing gene expression programs and RNA maturation pathways, including miRNA biogenesis through Drosha interaction. It plays roles in chromatin remodelling and epigenetic regulation that affect cellular differentiation and development. EWSR1 modulates mitochondrial biogenesis and energy homeostasis by controlling stability of PGC-1α, a critical regulator of mitochondrial. Mutations and fusion proteins involving EWSR1 are linked to cancers, neurological diseases such as Amyotrophic lateral sclerosis (ALS). EWSR1 fusion proteins drive certain sarcoma oncogenesis, however functions outside cancer remain broad but less well-defined (40). Importantly, EWSR1 acts as a host factor for Hepatitis C virus (HCV) by binding to a critical RNA element (CRE), modulating its activity, physically associating with viral replication sites, and thereby facilitating efficient viral RNA replication and production of infectious virus. This mechanism appears specific to replication rather than translation of viral proteins (41). Besides, EWSR1 acts as a host factor enhancing the replication of SARS-CoV-2 by binding and promoting the helicase function of NSP13 (33). On the contrary, our study shows that binding of EWSR1 to CHIKV nsP2 inhibits its helicase activity and reduces CHIKV replication.

Our studies show that EWSR1 binds both 3NCR48 and 5NCR-RC50 sequences, located in the plus- and minus-sense CHIKV genomic RNAs, respectively. Direct binding of these sequences to purified EWSR1 was shown by EMSA as well as BLI. The protein was also shown to bind the CHIKV RNA in the virus-infected cells. During the CHIKV infection, the EWSR1 protein was shown to accumulate in the cytoplasm. This is consistent with the fact that alphavirus RNA replication takes place in the cytoplasm (42, 43). During HCV infection also, EWSR1 was shown to relocate to the cytoplasm (41). EWSR1 is functionally hijacked by binding to SARS-CoV2 NSP13 helicase, and likely to be relocalized from the nucleus to the cytoplasm. The modulation of the EWSR1 protein level and its translocation during SARS-CoV2 replication, however, was not reported (33).

In the case of HCV as well as SARS-CoV2, EWSR1 was a positive modulator of the virus replication (33, 41). Our study showed that CHIKV replication was significantly enhanced in siRNA-mediated EWSR1-depleted HEK293T cells as well as in the EWSR1 KO HAP1 cells. On the other hand, ectopic expression of EWSR1 in HEK293T cells or in the EWSR1 KO HAP1 cells significantly suppressed CHIKV replication. These data demonstrate an antiviral role for EWSR1 in CHIKV replication.

There are several examples of host proteins that bind the NCRs of RNA viruses and have a proviral role. Thus, La Protein binds to the 3′-NCR of HCV RNA and promotes viral RNA stability and replication (44). Polypyrimidine tract binding protein (PTB) binds to the 5′-and 3′-NCRs of multiple RNA viruses such as Poliovirus and HCV. It enhances viral RNA translation and replication by facilitating RNA circularization (45). RPL17 and YBX1 proteins bind the HCV 3′-NCR to enhance production of infectious particles (46). NF90/NF110 binds viral 5′- and 3′-NCRs and stabilizes viral RNA, enhancing replication for Dengue virus and HCV (47, 48). Heterogeneous nuclear ribonucleoprotein hnRNP A1 binds to the NCRs of various viruses including SINV, EV71, HCV, facilitating viral RNA replication (16, 49). In case of Foot-and-mouth disease virus (FMDV), Poly(A)-binding protein (PABP) interacts with 3′-NCR and promotes translation initiation and viral RNA stability (50). Enolase (ENO), a host glycolytic enzyme, binds Bamboo mosaic virus (BaMV) NCRs to enhance replication (51). We previously showed that Nucleolin protein binds 3’-NCR of Japanese encephalitis virus (JEV) and enhances virus replication (52).

Importantly, there are also many examples of host proteins binding the NCRs of RNA viruses showing antiviral activity. The 5′-NCR of alphaviruses, like CHIKV, contains conserved RNA structures and a 5′-ppp moiety that binds IFIT1 protein, thereby blocking the assembly of the translation initiation complex. This inhibits viral protein synthesis, resulting in virus replication inhibition (53). For another alphavirus Venezuelan equine encephalitis virus, IFIT2 protein binds the 3’-NCR and modulates virus replication by a translation-independent mechanism (54). In case of SARS-CoV-2, RNA binding protein RBM24 was shown to bind the 5’-NCR, preventing 80S ribosome assembly, which in turn inhibits polyproteins translation and the virus replication (55). In another report, Lamp2 protein was shown to bind the 5’-NCR of the SARS-CoV-2 genome. Its level was upregulated in virus-infected cells; the absence of Lamp2a enhanced the viral RNA levels whereas its overexpression significantly reduced the viral RNA level (56). However, the mechanism of antiviral action was not studied. Yet another protein Pseudouridine synthase 4 (Pus4) was shown to bind specifically to the 3’-NCR of the Brome mosaic virus RNA (57). Here, the protein binding interfered with the plus-sense genomic RNA synthesis reducing the virus replication. This is similar to the present report showing specific binding of EWSR1 to the 3’-NCR of the plus-sense CHIKV genomic RNA and the 3’-end of the replication intermediate RNA, reduced plus- and minus-sense genomic RNA synthesis in the presence of overexpressed EWSR1, and reduced virus replication.

Our studies have shown that both CHIKV nsP2 and EWSR1 bind the 3NCR48 sequence. CHIKV nsP2 helicase and the replicase complex have to move along the genomic RNA template for its unwinding and copying. EWSR1 binding to 3NCR48 could sterically hinder or physically block CHIKV nsP2 and the viral replicase complex access and assembly, or movement, at the 3′-end of the CHIKV plus-sense genomic RNA, thus inhibiting the minus-sense RNA synthesis. Similarly, EWSR1 binding to 5NCR-RC50 sequence could obstruct the access and movement of CHIKV nsP2 and the viral replicase at the 3′-end of the replication-intermediate minus-sense CHIKV RNA, inhibiting the synthesis of the plus-sense CHIKV genomic RNA, resulting in the inhibition of the virus replication. We have also shown EWSR1 binding to CHIKV nsP2 resulting in the inhibition of its helicase activity. EWSR1 at the viral RNA replication site may bind CHIKV nsP2 making it unavailable for the RNA replication, resulting in the inhibition of the viral RNA synthesis and reduced virus titers. This presents an alternate mechanism of the EWSR1-mediated antiviral effect seen during the CHIKV replication.

The 3’-NCR’s 48-nucleotide conserved sequence represented in the 3NCR48 RNA has been predicted to form a hairpin secondary structure (15, 58, 59). EWSR1 binding may stabilize or destabilize such key RNA elements, altering local secondary or tertiary RNA structures and thereby affecting the efficiency of replication. EWSR1 could help recruit host factors involved in RNA decay or restrict viral RNA from protective compartments, leading to enhanced viral RNA degradation. Role of these mechanism in EWSR1-mediated inhibition of CHIKV replication, if any, needs further investigations.

## Data Availability Statement

All the data related to this work are included in the manuscript.

## Competing interest declaration

The authors declare no competing interests.

## Acknowledgements

The work was supported by grant no. JCB/2021/000015 from the Science and Engineering Research Board, Govt. of India, and grant no. BT/PR54847/BMS2/156/114/2024 from the Department of Biotechnology, Govt. of India to SV. SB received the Senior Research Fellowship from the Department of Biotechnology, Govt. of India. We thank Dr. Ramandeep Singh for the PPK-1 protein. We acknowledge the expert advice of Dr. Nirpendra Singh, Dr. Tushar Kanti Maiti, Dr. Amulya Panda, Dharmendra Gupta, and the use of the Advanced technology Platform Centre (ATPC) facilities at Regional Centre for Biotechnology.

## REFERENCES

1. Freppel W, Silva LA, Stapleford KA, Herrero LJ. 2024. Pathogenicity and virulence of chikungunya virus. Virulence 15:2396484.

2. Anonymous. https://www.ecdc.europa.eu/en/chikungunya-monthly. Accessed

3. Ly H. 2024. Ixchiq (VLA1553): The first FDA-approved vaccine to prevent disease caused by Chikungunya virus infection. Virulence 15:2301573.

4. Tindale LC, Richardson JS, Anderson DM, Mendy J, Muhammad S, Loreth T, Tredo SR, Ramanathan R, Jenkins VA, Bedell L, Ajiboye P, Group E-C-S. 2025. Chikungunya virus virus-like particle vaccine safety and immunogenicity in adults older than 65 years: a phase 3, randomised, double-blind, placebo-controlled trial. Lancet 405:1353–1361.

5. Richardson JS, Anderson DM, Mendy J, Tindale LC, Muhammad S, Loreth T, Tredo SR, Warfield KL, Ramanathan R, Caso JT, Jenkins VA, Ajiboye P, Bedell L, Group E-C-S. 2025. Chikungunya virus virus-like particle vaccine safety and immunogenicity in adolescents and adults in the USA: a phase 3, randomised, double-blind, placebo-controlled trial. Lancet 405:1343–1352.

6. Strauss JH, Strauss EG. 1994. The alphaviruses: gene expression, replication, and evolution. Microbiol Rev 58:491–562.

7. Pletnev SV, Zhang W, Mukhopadhyay S, Fisher BR, Hernandez R, Brown DT, Baker TS, Rossmann MG, Kuhn RJ. 2001. Locations of carbohydrate sites on alphavirus glycoproteins show that E1 forms an icosahedral scaffold. Cell 105:127–136.

8. Barraza Sanchez JJ, Volpe S, Faraj SE, Filomatori CV. 2025. Functional RNA Elements in Alphavirus Genomes. Curr Microbiol 82:364.

9. Kulasegaran-Shylini R, Atasheva S, Gorenstein DG, Frolov I. 2009. Structural and functional elements of the promoter encoded by the 5’ untranslated region of the Venezuelan equine encephalitis virus genome. J Virol 83:8327–39.

10. Frolov I, Hardy R, Rice CM. 2001. Cis-acting RNA elements at the 5’ end of Sindbis virus genome RNA regulate minus- and plus-strand RNA synthesis. RNA 7:1638–51.

11. Hyde JL, Chen R, Trobaugh DW, Diamond MS, Weaver SC, Klimstra WB, Wilusz J. 2015. The 5’ and 3’ ends of alphavirus RNAs--Non-coding is not non-functional. Virus Res 206:99–107.

12. Rupp JC, Sokoloski KJ, Gebhart NN, Hardy RW. 2015. Alphavirus RNA synthesis and non-structural protein functions. J Gen Virol 96:2483–2500.

13. Morley VJ, Noval MG, Chen R, Weaver SC, Vignuzzi M, Stapleford KA, Turner PE. 2018. Chikungunya virus evolution following a large 3’UTR deletion results in host-specific molecular changes in protein-coding regions. Virus Evol 4:vey012.

14. Garcia-Moreno M, Sanz MA, Carrasco L. 2016. A Viral mRNA Motif at the 3’-Untranslated Region that Confers Translatability in a Cell-Specific Manner. Implications for Virus Evolution. Sci Rep 6:19217.

15. Bardossy ES, Volpe S, Alvarez DE, Filomatori CV. 2023. A conserved Y-shaped RNA structure in the 3’UTR of chikungunya virus genome as a host-specialized element that modulates viral replication and evolution. PLoS Pathog 19:e1011352.

16. Lin JY, Shih SR, Pan M, Li C, Lue CF, Stollar V, Li ML. 2009. hnRNP A1 interacts with the 5’ untranslated regions of enterovirus 71 and Sindbis virus RNA and is required for viral replication. J Virol 83:6106–14.

17. George J, Raju R. 2000. Alphavirus RNA genome repair and evolution: molecular characterization of infectious sindbis virus isolates lacking a known conserved motif at the 3’ end of the genome. J Virol 74:9776–85.

18. Dickson AM, Anderson JR, Barnhart MD, Sokoloski KJ, Oko L, Opyrchal M, Galanis E, Wilusz CJ, Morrison TE, Wilusz J. 2012. Dephosphorylation of HuR protein during alphavirus infection is associated with HuR relocalization to the cytoplasm. J Biol Chem 287:36229–38.

19. Pardigon N, Strauss JH. 1996. Mosquito homolog of the La autoantigen binds to Sindbis virus RNA. J Virol 70:1173–81.

20. Ta M, Vrati S. 2000. Mov34 protein from mouse brain interacts with the 3’ noncoding region of Japanese encephalitis virus. J Virol 74:5108–15.

21. Singh SM, Panda AK. 2005. Solubilization and refolding of bacterial inclusion body proteins. J Biosci Bioeng 99:303–10.

22. Sharma KB, Subramani C, Ganesh K, Sharma A, Basu B, Balyan S, Sharma G, Ka S, Deb A, Srivastava M, Chugh S, Sehrawat S, Bharadwaj K, Rout A, Sahoo PK, Saurav S, Motiani RK, Singh R, Jain D, Asthana S, Wadhwa R, Vrati S. 2024. Withaferin A inhibits Chikungunya virus nsP2 protease and shows antiviral activity in the cell culture and mouse model of virus infection. PLoS Pathog 20:e1012816.

23. Law YS, Wang S, Tan YB, Shih O, Utt A, Goh WY, Lian BJ, Chen MW, Jeng US, Merits A, Luo D. 2021. Interdomain Flexibility of Chikungunya Virus nsP2 Helicase-Protease Differentially Influences Viral RNA Replication and Infectivity. J Virol 95.

24. Das PK, Merits A, Lulla A. 2014. Functional cross-talk between distant domains of chikungunya virus non-structural protein 2 is decisive for its RNA-modulating activity. J Biol Chem 289:5635–53.

25. Singh H, Mudgal R, Narwal M, Kaur R, Singh VA, Malik A, Chaudhary M, Tomar S. 2018. Chikungunya virus inhibition by peptidomimetic inhibitors targeting virus-specific cysteine protease. Biochimie 149:51–61.

26. Saha A, Acharya BN, Priya R, Tripathi NK, Shrivastava A, Rao MK, Kesari P, Narwal M, Tomar S, Bhagyawant SS, Parida M, Dash PK. 2018. Development of nsP2 protease based cell free high throughput screening assay for evaluation of inhibitors against emerging Chikungunya virus. Sci Rep 8:10831.

27. Saisawang C, Sillapee P, Sinsirimongkol K, Ubol S, Smith DR, Ketterman AJ. 2015. Full length and protease domain activity of chikungunya virus nsP2 differ from other alphavirus nsP2 proteases in recognition of small peptide substrates. Biosci Rep 35.

28. Khan AH, Morita K, Parquet MDC, Hasebe F, Mathenge EGM, Igarashi A. 2002. Complete nucleotide sequence of chikungunya virus and evidence for an internal polyadenylation site. J Gen Virol 83:3075–3084.

29. Sun K, Appadoo F, Liu Y, Muller M, Macfarlane C, Harris M, Tuplin A. 2024. A novel interaction between the 5’ untranslated region of the Chikungunya virus genome and Musashi RNA binding protein is essential for efficient virus genome replication. Nucleic Acids Res 52:10654–10667.

30. Gorchakov R, Hardy R, Rice CM, Frolov I. 2004. Selection of functional 5’ cis-acting elements promoting efficient sindbis virus genome replication. J Virol 78:61–75.

31. Zakaryan RP, Gehring H. 2006. Identification and characterization of the nuclear localization/retention signal in the EWS proto-oncoprotein. J Mol Biol 363:27–38.

32. Law YS, Utt A, Tan YB, Zheng J, Wang S, Chen MW, Griffin PR, Merits A, Luo D. 2019. Structural insights into RNA recognition by the Chikungunya virus nsP2 helicase. Proc Natl Acad Sci U S A 116:9558–9567.

33. Zeng H, Gao X, Xu G, Zhang S, Cheng L, Xiao T, Zu W, Zhang Z. 2022. SARS-CoV-2 helicase NSP13 hijacks the host protein EWSR1 to promote viral replication by enhancing RNA unwinding activity. Infect Med (Beijing) 1:7–16.

34. Pieterse L, McDonald M, Abraham R, Griffin DE. 2024. Heterogeneous Ribonucleoprotein K Is a Host Regulatory Factor of Chikungunya Virus Replication in Astrocytes. Viruses 16.

35. Ander SE, Carpentier KS, Sanders W, Lucas CJ, Jolly AJ, Johnson CN, Hawman DW, Heise MT, Moorman NJ, Morrison TE. 2024. A 44-Nucleotide Region in the Chikungunya Virus 3’ UTR Dictates Viral Fitness in Disparate Host Cells. Viruses 16.

36. Garneau NL, Sokoloski KJ, Opyrchal M, Neff CP, Wilusz CJ, Wilusz J. 2008. The 3’ untranslated region of sindbis virus represses deadenylation of viral transcripts in mosquito and Mammalian cells. J Virol 82:880–92.

37. Pardigon N, Strauss JH. 1992. Cellular proteins bind to the 3’ end of Sindbis virus minus-strand RNA. J Virol 66:1007–15.

38. Pardigon N, Lenches E, Strauss JH. 1993. Multiple binding sites for cellular proteins in the 3’ end of Sindbis alphavirus minus-sense RNA. J Virol 67:5003–11.

39. Fata CL, Sawicki SG, Sawicki DL. 2002. Alphavirus minus-strand RNA synthesis: identification of a role for Arg183 of the nsP4 polymerase. J Virol 76:8632–40.

40. Lee J, Nguyen PT, Shim HS, Hyeon SJ, Im H, Choi MH, Chung S, Kowall NW, Lee SB, Ryu H. 2019. EWSR1, a multifunctional protein, regulates cellular function and aging via genetic and epigenetic pathways. Biochim Biophys Acta Mol Basis Dis 1865:1938–1945.

41. Oakland TE, Haselton KJ, Randall G. 2013. EWSR1 binds the hepatitis C virus cis-acting replication element and is required for efficient viral replication. J Virol 87:6625–34.

42. Spuul P, Balistreri G, Hellstrom K, Golubtsov AV, Jokitalo E, Ahola T. 2011. Assembly of alphavirus replication complexes from RNA and protein components in a novel trans-replication system in mammalian cells. J Virol 85:4739–51.

43. Froshauer S, Kartenbeck J, Helenius A. 1988. Alphavirus RNA replicase is located on the cytoplasmic surface of endosomes and lysosomes. J Cell Biol 107:2075–86.

44. Spangberg K, Wiklund L, Schwartz S. 2001. Binding of the La autoantigen to the hepatitis C virus 3’ untranslated region protects the RNA from rapid degradation in vitro. J Gen Virol 82:113–120.

45. Gutierrez AL, Denova-Ocampo M, Racaniello VR, del Angel RM. 1997. Attenuating mutations in the poliovirus 5’ untranslated region alter its interaction with polypyrimidine tract-binding protein. J Virol 71:3826–33.

46. Liu J, Ito M, Liu L, Nakashima K, Satoh S, Konno A, Suzuki T. 2024. Involvement of ribosomal protein L17 and Y-box binding protein 1 in the assembly of hepatitis C virus potentially via their interaction with the 3’ untranslated region of the viral genome. J Virol 98:e0052224.

47. Gomila RC, Martin GW, Gehrke L. 2011. NF90 binds the dengue virus RNA 3’ terminus and is a positive regulator of dengue virus replication. PLoS One 6:e16687.

48. Isken O, Baroth M, Grassmann CW, Weinlich S, Ostareck DH, Ostareck-Lederer A, Behrens SE. 2007. Nuclear factors are involved in hepatitis C virus RNA replication. RNA 13:1675–92.

49. Kim CS, Seol SK, Song OK, Park JH, Jang SK. 2007. An RNA-binding protein, hnRNP A1, and a scaffold protein, septin 6, facilitate hepatitis C virus replication. J Virol 81:3852–65.

50. Rodriguez Pulido M, Serrano P, Saiz M, Martinez-Salas E. 2007. Foot-and-mouth disease virus infection induces proteolytic cleavage of PTB, eIF3a,b, and PABP RNA-binding proteins. Virology 364:466–74.

51. Lin KY, Huang YW, Hou LY, Chen HC, Wu Y, Chen IH, Huang YP, Lee SC, Hu CC, Tsai CH, Hsu YH, Lin NS. 2025. Proviral insights of glycolytic enolase in Bamboo mosaic virus replication associated with chloroplasts and mitochondria. Proc Natl Acad Sci U S A 122:e2415089122.

52. Deb A, Nagpal S, Yadav RK, Thakur H, Nair D, Krishnan V, Vrati S. 2024. Japanese encephalitis virus NS5 protein interacts with nucleolin to enhance the virus replication. J Virol 98:e0085824.

53. Reynaud JM, Kim DY, Atasheva S, Rasalouskaya A, White JP, Diamond MS, Weaver SC, Frolova EI, Frolov I. 2015. IFIT1 Differentially Interferes with Translation and Replication of Alphavirus Genomes and Promotes Induction of Type I Interferon. PLoS Pathog 11:e1004863.

54. Hickson SE, Brekke E, Schwerk J, Saluhke I, Zaver S, Woodward J, Savan R, Hyde JL. 2025. Sequence Diversity in the 3’ Untranslated Region of Alphavirus Modulates IFIT2-Dependent Restriction in a Cell Type-Dependent Manner. J Interferon Cytokine Res 45:133–149.

55. Yao Y, Sun H, Chen Y, Tian L, Huang D, Liu C, Zhou Y, Wang Y, Wen Z, Yang B, Chen X, Pei R. 2023. RBM24 inhibits the translation of SARS-CoV-2 polyproteins by targeting the 5’-untranslated region. Antiviral Res 209:105478.

56. Verma R, Saha S, Kumar S, Mani S, Maiti TK, Surjit M. 2021. RNA-Protein Interaction Analysis of SARS-CoV-2 5’ and 3’ Untranslated Regions Reveals a Role of Lysosome-Associated Membrane Protein-2a during Viral Infection. mSystems 6:e0064321.

57. Zhu J, Gopinath K, Murali A, Yi G, Hayward SD, Zhu H, Kao C. 2007. RNA-binding proteins that inhibit RNA virus infection. Proc Natl Acad Sci U S A 104:3129–34.

58. Madden EA, Plante KS, Morrison CR, Kutchko KM, Sanders W, Long KM, Taft-Benz S, Cruz Cisneros MC, White AM, Sarkar S, Reynolds G, Vincent HA, Laederach A, Moorman NJ, Heise MT. 2020. Using SHAPE-MaP To Model RNA Secondary Structure and Identify 3’UTR Variation in Chikungunya Virus. J Virol 94.

59. Schneider AB, Ochsenreiter R, Hostager R, Hofacker IL, Janies D, Wolfinger MT. 2019. Updated Phylogeny of Chikungunya Virus Suggests Lineage-Specific RNA Architecture. Viruses 11.

